# DamID transcriptional profiling identifies the Snail/Scratch transcription factor Kahuli as Alk target in the *Drosophila* visceral mesoderm

**DOI:** 10.1101/2021.01.25.428051

**Authors:** Patricia Mendoza-Garcia, Swaraj Basu, Sanjay Kumar Sukumar, Badrul Arefin, Georg Wolfstetter, Vimala Anthonydhason, Linnea Molander, Henrik Lindehell, Jan Larsson, Erik Larsson, Mats Bemark, Ruth H. Palmer

## Abstract

Development of the midgut visceral muscle of *Drosophila* crucially depends on Anaplastic Lymphoma Kinase (Alk) receptor tyrosine kinase (RTK) signalling, which is needed to specify founder cells (FCs) in the circular visceral mesoderm (VM). While activation of the Alk receptor by its ligand Jelly Belly (Jeb) is well characterized, only a small number of target molecules have been identified. Here, we assayed RNA polymerase II (Pol II) occupancy in VM cells by using the targeted DamID (TaDa) approach. To identify Alk targets we employed comparative analysis of embryos overexpressing Jeb *versus* embryos with abrogated Alk activity, revealing differential expression of a number of genes, including the Snail/Scratch family transcription factor *Kahuli* (*Kah*). Upon further *in vivo* validation, we confirmed that Alk signalling regulates *Kah* mRNA expression in the VM. We show that *Kah* mutants display defects in the formation of midgut constrictions, similar to that of *pointed* (*pnt*) mutants. Analysis of publicly available ChIP data defined a Kah target-binding site similar to that of Snail. In addition, we compared genes that were differentially expressed in *Kah* mutants with publicly available Kah- and Pnt-ChIP datasets identifying a set of common target genes putatively regulated by Kah and Pnt in midgut constriction. Taken together, we (i) report a rich dataset of Alk responsive loci in the embryonic VM, (ii) provide the first functional characterization of the Kah transcription factor, identifying a role in embryonic midgut constriction, and (iii) suggest a model in which Kah and Pnt cooperate in embryonic midgut morphogenesis.

## Introduction

Receptor tyrosine kinase (RTK) signalling enables transduction of extracellular signals into the cell and is essential in a wide range of developmental processes such as cell fate determination, differentiation, patterning, proliferation, growth, and survival. Ligand-dependent RTK activation is conserved among metazoans, leading to engagement of signal transduction adaptor proteins, serine/threonine kinases, and transcription factors that regulate gene expression and promote a wide range of intracellular responses. In *Drosophila melanogaster*, the Anaplastic Lymphoma Kinase (Alk) RTK and its ligand Jelly Belly (Jeb), are involved in the development of the visceral mesoderm (VM) where they drive a signalling pathway required for specification of muscle founder cells (FCs) (Englund et al., 2003; Lee et al., 2003; Stute et al., 2004). Ligand-stimulated activation of Alk acts through the guanosine triphosphatase Ras, the scaffold protein connector enhancer of kinase suppressor of Ras (Cnk) and Aveugle/Hyphen (Ave/Hyp) to drive the mitogen-activated protein kinase/extracellular signal-regulated kinase (MAPK/ERK) pathway via activation of the serine-threonine kinases Raf and MEK (Englund et al., 2003; Lee et al., 2003; Stute et al., 2004; Wolfstetter et al., 2017).

During *Drosophila* development, the post-gastrulation mesoderm is partitioned along the dorso-ventral axis due to inductive inputs from the ectoderm such as Decapentaplegic (Dpp), which induces high levels of Tinman (Tin) and subsequently Bagpipe (Bap), leading to specification of the VM (Frasch, 1995). This early VM consists of naïve Alk expressing myoblasts that are specified to become either founder cells (FCs) or fusion competent myoblasts (FCMs). Specification of FCs requires activation of the Alk signal transduction cascade by Jeb secreted from the adjacent somatic mesoderm (Englund et al., 2003; Lee et al., 2003; Stute et al., 2004). After specification, FCs fuse with FCMs to form binucleate myotubes (Campos-Ortega, 1997; Klapper et al., 2002; Lee et al., 2006; Martin et al., 2001; Poulson, 1950). Fusion of FCs and FCMs is required for the formation of circular visceral muscles, upon which the longitudinal muscle precursors migrate, and ultimately form a web of interconnected muscles that surround the midgut endoderm (Georgias et al., 1997; Klapper et al., 2002; Kusch and Reuter, 1999; Martin et al., 2001; Rudolf et al., 2014). Although Alk is expressed throughout the VM, only the ventral-most row of cells within each cluster are exposed to Jeb (Englund et al., 2003; Lee et al., 2003; Loren et al., 2001; Stute et al., 2004). Alk signalling regulates transcription of FC-specific genes including *Hand, optomotor-blind-related-gene-1* (*org-1*), and *kin of irre* (*kirre*, also known as *dumbfounded - duf*) (Englund et al., 2003; Lee et al., 2003; Stute et al., 2004; Varshney and Palmer, 2006). Animals devoid of FCs specified by Jeb/Alk signalling do not undergo myoblast fusion and the visceral musculature fails to develop (Englund et al., 2003; Lee et al., 2003; Loren et al., 2001; Stute et al., 2004). The influence of Alk signalling on later events in visceral myogenesis is largely unclear, however Alk activity is required for *dpp* expression in the VM, and Alk mutants lack *dpp* and resultant pMAD signalling in both the VM and adjacent endoderm (Shirinian et al., 2007).

Although Alk signalling has been widely studied during *Drosophila* embryo development, assaying transcriptional effects specifically in the VM is challenging using traditional transcriptomics, and only a small number of Alk transcriptional targets have been described. To address this, we used Targeted DamID (TaDa) to determine genome-wide Alk regulated transcriptional events in the embryonic VM. TaDa exploits the activity of bacterial DNA adenine methyltransferase (Dam) fused to any protein of interest to allow determination of cell type-specific DNA-binding profiles, and has previously been used with RNA polymerases, transcription factors (TFs), and histone modifiers (e.g. histone deacetylase), among others (Aughey and Southall, 2016). TaDa can further be refined to address DNA-binding profiles in specific developmental tissues and at time points of interest using the well-established GAL4/UAS expression system (Brand and Perrimon, 1993; Southall et al., 2013). This tissue specific approach revealed known targets of Alk signalling in the VM, as well a large number of previously unidentified Alk target genes. Among these, we identified and validated the Snail/Scratch family transcription factor Kahuli (Kah) as a novel target of Jeb/Alk signalling in the VM. Loss of Alk signalling (in an *Alk* or *jeb* null mutant background) resulted in reduced *Kah* mRNA expression in FCs, while ectopic activation of Alk increased *Kah* expression. To gain further insight on *Kah* function, we generated and characterized *Kah* loss of function mutants, which specify FCs, but fail to form the first midgut constriction at later stages of embryonic development. We show that this defect in *Kah* mutants is similar to that previously described for *pnt* mutants, suggesting that Kah and Pnt function together to regulate this process. Combination of publicly available ChIP datasets for Kah and Pnt revealed a number of common targets, reinforcing the hypothesis that Kah and Pnt work together in midgut morphogenesis. Thus, our Alk activity dependent DamID approach successfully identified a number of Alk regulated transcriptional targets in the embryonic VM, including the Kah TF as an Alk target that is required for correct midgut constriction.

## Results

### Targeted DamID-derived transcriptional landscape of the *Drosophila* VM

To characterize Alk regulated transcriptional activity *in vivo,* we employed TaDa in the embryonic VM in which we genetically manipulated Alk signalling output. We used transgenic *Drosophila* strains expressing *Dam* methylase fused to *RNA-Pol II* transgene (here named *Dam-Pol II*) (Southall et al., 2013). Expression of *Dam-Pol II* was driven either generally in the mesoderm (*twi2xPE-GAL4*) or more specifically in the VM (*bap-GAL4*), resulting in methylation at GATC sites in the targeted tissue (Fig. 1A). To manipulate Alk signalling we used combinatorial expression of either *UAS-Jeb* which leads to ectopic activation of Alk or *UAS-Alk.EC.MYC* encoding the extracellular domain (ECD) of Alk that inhibits Alk signalling in a dominant-negative manner (further referred to as *UAS-Alk.DN*) (Bazigou et al., 2007). (Fig. 1A, C-E). Animals expressing *Dam-Pol II* alone in a wild-type background were employed to control for basal Dam-Pol II signal (Fig. 1A, C). Stage 10-13 embryos were collected, representing a developmental window during which Alk is activated in future visceral FCs, and experimental sampling was performed in triplicates. Methylated DNA from collected embryos was isolated and digested with the methylation specific Dpn I restriction endonuclease, followed by next-generation sequencing to identify genes that are transcriptional targets of Alk signalling (Fig. 1B). Our analysis of this dataset was based on previous pipelines developed for DamID (Maksimov et al., 2016; Tosti et al., 2018). The number of quality reads obtained were comparable between samples and replicates (above 15M reads per sample, Fig. S1A). After alignment to the *Drosophila* genome, sequencing depth was above 60% for every sample, with exception of one sample (summarized in Fig. S1B). A high degree of reproducibility was observed between biological replicates overexpressing Alk.DN and Dam-Pol II. In contrast, replicates of samples ectopically expressing Jeb displayed substantial variation (Fig. S1C). Therefore, we employed *twi2xPE-GAL4* and *bap-Gal4 driven UAS-jeb* NGS data for qualitative analyses only.

**Figure 1.**
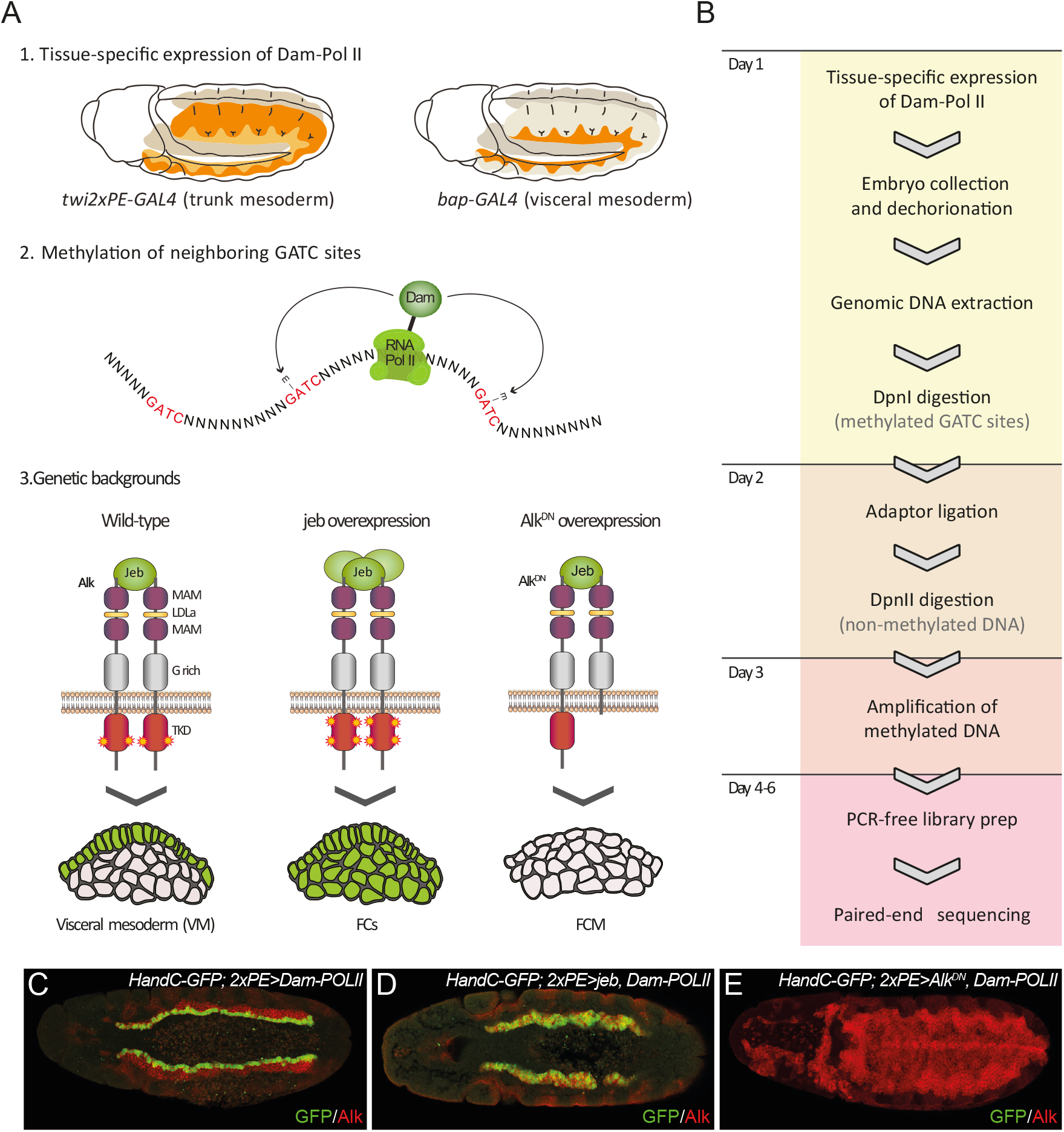
TaDa allows *in vivo* VM-specific transcriptional profiling. **A.** Outline of experimental approach: the lineage-specific GAL4 drivers *bap-GAL4* and *twi.2xPE-GAL4* were used to drive expression of Dam-Pol II (1), leading to methylation of GATC sequences in the genome (2), analysis was performed under wild-type Alk signalling, ligand (*jeb*) overexpression (Alk signalling activation) or dominant negative inhibition of Alk signalling (*Alk.DN*) (3). **B.** Experimental flow chart from targeted DamID expression to library preparation and sequencing. **C-E.** *HandC-GFP* reporter gene expression in the three genetic backgrounds included in our TaDa analysis: GFP labels the ventral-most FC row in embryos with wild-type Alk signalling (C), all VM cells upon *twi2xPE-Gal4* driven *jeb* overexpression (D), or is non-detectable in *twi2xPE-Gal4>Alk^DN^* embryos (E).

To assess whether our DamID approach recapitulates transcriptionally active regions of the genome, we performed a meta-analysis of Dam-Pol II occupancy, as indicated by GATC associated reads (see Materials and Methods for details), relative to the distance to the closest transcription start site (TSS). When comparing all GATC motifs (Non Dam-Pol II) and random regions on the genome to Dam-Pol II methylated GATC sites (Dam-Pol II) we observed a tendency for methylated GATC sites to accumulate close to TSSs (Fig. S2A). In addition, we compared our DamID results with previously published RNA-seq data from isolated mesoderm cells (NCBI BioProject, accession number PRJEB11879). In agreement with our previous observations, the Dam-Pol II binding profile along all annotated genes is consistent with an RNA expression profile of mesodermal cells (Fig. S2B-C), demonstrating that the Dam-Pol II binding in our analyses reflects Pol II *in vivo* occupancy.

### TaDa identifies Alk-regulated loci in the *Drosophila* VM

To detect differential gene expression between *Dam-Pol II* and *Alk.DN* samples, we clustered neighboring GATC associated reads, maximum 350 bp apart (median GATC fragment distance for the *Drosophila* genome) into peaks (Tosti et al., 2018). Most peaks were associated with a single GATC (Fig. 2A). We then calculated the mean fold change ratios for all GATCs falling into each peak across annotated transcripts (*Alk.DN/Dam-Pol II*), and a false discovery rate (FDR) was assigned to each peak. Each gene along the genome was assigned an overlapping peak with the minimum FDR value, and its logFC and FDR were used for differential expression analysis visualization on a volcano plot (Fig. 2B).

**Figure 2.**
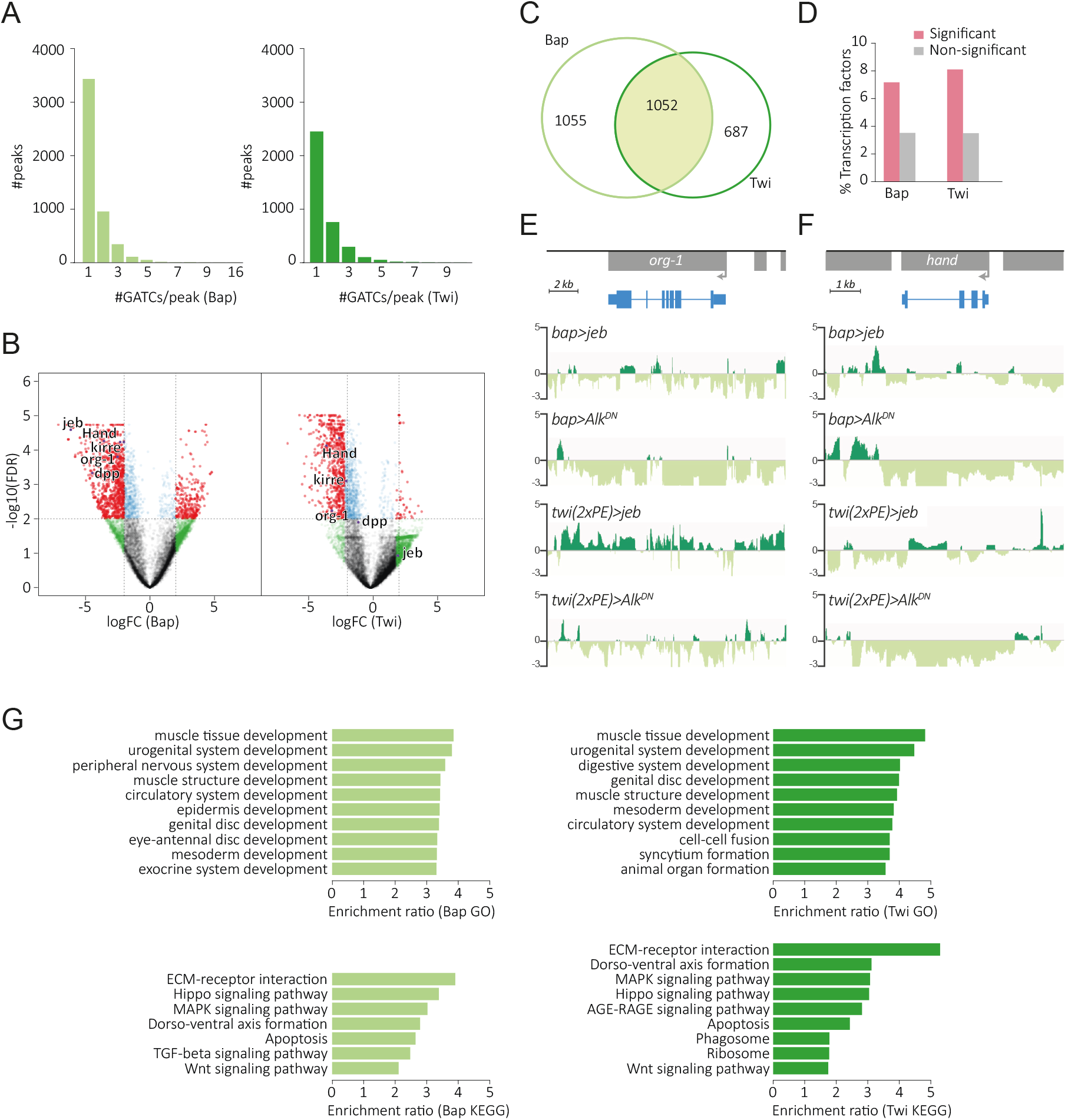
Significant peaks and associated genes identified by TaDa. LogFC of reads mapped to GATCs obtained by comparing *UAS-Alk.DN* samples against Dam-Pol II samples separately for *bap-GAL4* (Bap) and *twi.2xPE-GAL4* (Twi) samples. Peaks were built by clustering GATC sites at median GATC fragment distance for the *Drosophila* genome. The logFC represents the mean logFC of all GATCs falling inside the peak. **A.** Distribution of peaks formed by clustering, expressed as number of GATC sites per peak. **B.** Each gene was assigned an overlapping peak with a minimum FDR value, and both logFC and FDR for the assigned peaks are shown as a volcano plot. **C.** Number of genes associated with peaks at an FDR < 0.01 for Bap and Twi datasets. **D.** Genes associated with Bap and Twi peaks (FDR < 0.01) are enriched for transcription factors, compared to the remaining set of genes in both instances (Fisher test, p < 2e-16). **E, F.** Differential Dam-Pol II occupancy over *org-1* (E) and *Hand* (F) (known Alk targets) loci upon *jeb* or *Alk.DN* overexpression. Scale bars indicate log2FC between *UAS*-Dam-Pol II (reference) and *UAS-Dam-Pol II, UAS-jeb* or *UAS-Dam-Pol II, UAS-Alk.DN* samples. **G.** Enrichment of GO terms and KEGG pathways (FDR < 0.05) for the genes associated with significant peaks (Bap and Twi).

For statistical analyses, an FDR < 0.01 (log10 (FDR) > 2) was considered significant. In total, we identified significant change in Dam-Pol II occupancy on 1739 genes in the *twi.2xPE-GAL4* samples (Twi) and 2107 genes in the *bap-GAL4* samples (Bap), with an overlap of 1052 genes between samples (Fig. 2C). The identified genes included known targets of Alk signalling such as *Hand, org-1, kirre* and *dpp* (Fig. 2B, E-F) (Lee et al., 2003; Loren et al., 2003; Shirinian et al., 2007; Stute et al., 2004; Varshney and Palmer, 2006), demonstrating that the TaDa approach was successfully able to identify Alk targets in the VM.

Alk signalling in the VM is known to control FC specification, therefore we expected transcriptional activation of factors involved in this process. With TaDa, we observed peak enrichment for transcription factors in both Bap and Twi datasets, including *Hand* and *org-1* (Fig. 2B, D). We also observed genes known to play a role in VM cell fusion, such as *kirre* (up-regulated), *sns* and *Vrp1* (down-regulated) (Fig. 2B). Moreover, we identified factors involved in signalling pathways known to be active during development of the mesoderm, musculature and nervous system (Fig. 2G).

Qualitative analysis of peak-associated genes with the lowest p-values showed differential Dam-Pol II occupancy between Jeb and Alk.DN overexpression samples. At the individual gene level, occupancy of Dam-Pol II reveals similar binding profiles for *twi.2xPE-GAL4* and *bap-GAL4* samples (Fig. 3A-F). Further *in vivo* validation of a selection of highly significant genes differentially expressed by *in situ* hybridization showed them to be actively expressed in the VM. These included the *Kahuli* (*Kah*) TF, the transmembrane protein *failed axon connections* (*fax*), the *PAR-domain protein 1 (Pdp1), CG11658, CG5149*, and the SUMO family protein *smt3* (Fig. 3A-F). Taken together, our bioinformatics analysis and experimental validation shows the effectiveness of TaDa in the identification of novel transcriptional regulation events downstream of Alk in the embryonic VM.

**Figure 3.**
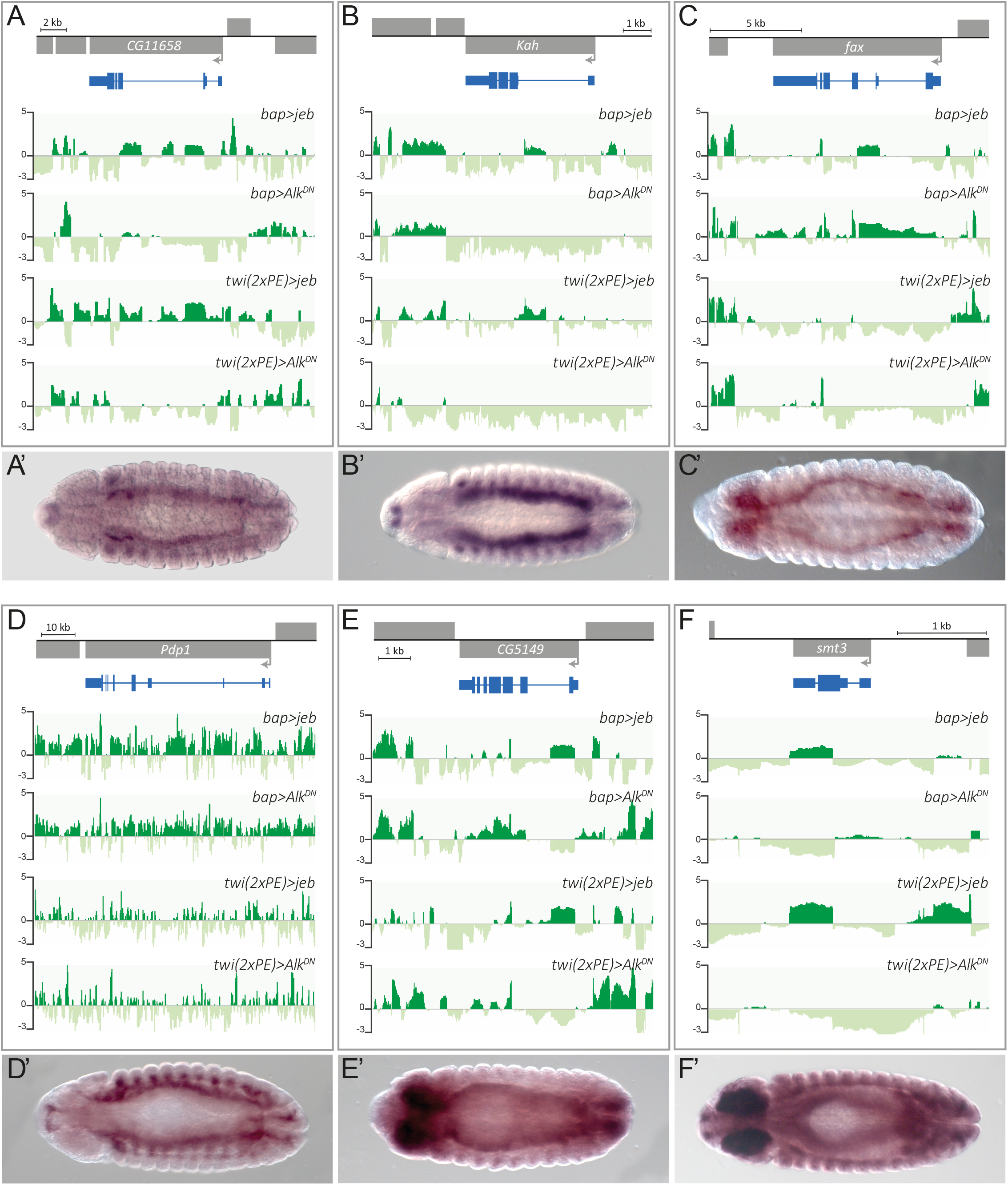
Validation of selected TaDa identified gene expression in the VM. **A-F.** Dam-Pol II occupancy of selected candidate loci using *bap-* and *twi.2xPE-GAL4* drivers. Scale bars represent logFC between *UAS-Dam-Pol II* (reference) and *UAS-Dam-Pol II*, *UAS-jeb* or *UAS-Dam-Pol* II, *UAS-Alk.DN* samples. **A’-F’.** Expression patterns of the respective candidate genes at stage 13.

### Alk targets identified by TaDa are enriched in the visceral mesoderm

To further validate the transcriptional activation of the identified candidates upon Alk signalling we employed single-cell RNA-sequencing (scRNA-seq) profiling on cells isolated from dissociated stage 10-13 embryos. We used live/dead cell markers to isolate living cells with flow cytometry (Fig. 4A). After quality control filtering in the Seurat R toolkit, a total of 1055 cells from wild-type embryos were further analyzed (Satija et al., 2015). Unsupervised clustering of cells based on gene expression profiles identified 13 cell clusters with distinct transcriptional profiles that could be assigned to distinct cell lineages (Fig. 4B, C). When visualized in a two-dimensional uniform manifold approximation and projection (UMAP) plot the clusters distributed into 4 main groups, one of which comprised clusters of mesodermal origin (1, 2, 8, 9 and 10) (Fig. 4B, C). Within this group, the cluster representing the VM was identified by plotting combinatorial gene expression of known factors involved in VM development, such as *biniou (bin), bagpipe (bap), org-1, Hand,* and *Fasciclin 3 (Fas3*) (Fig. 4D).

**Figure 4.**
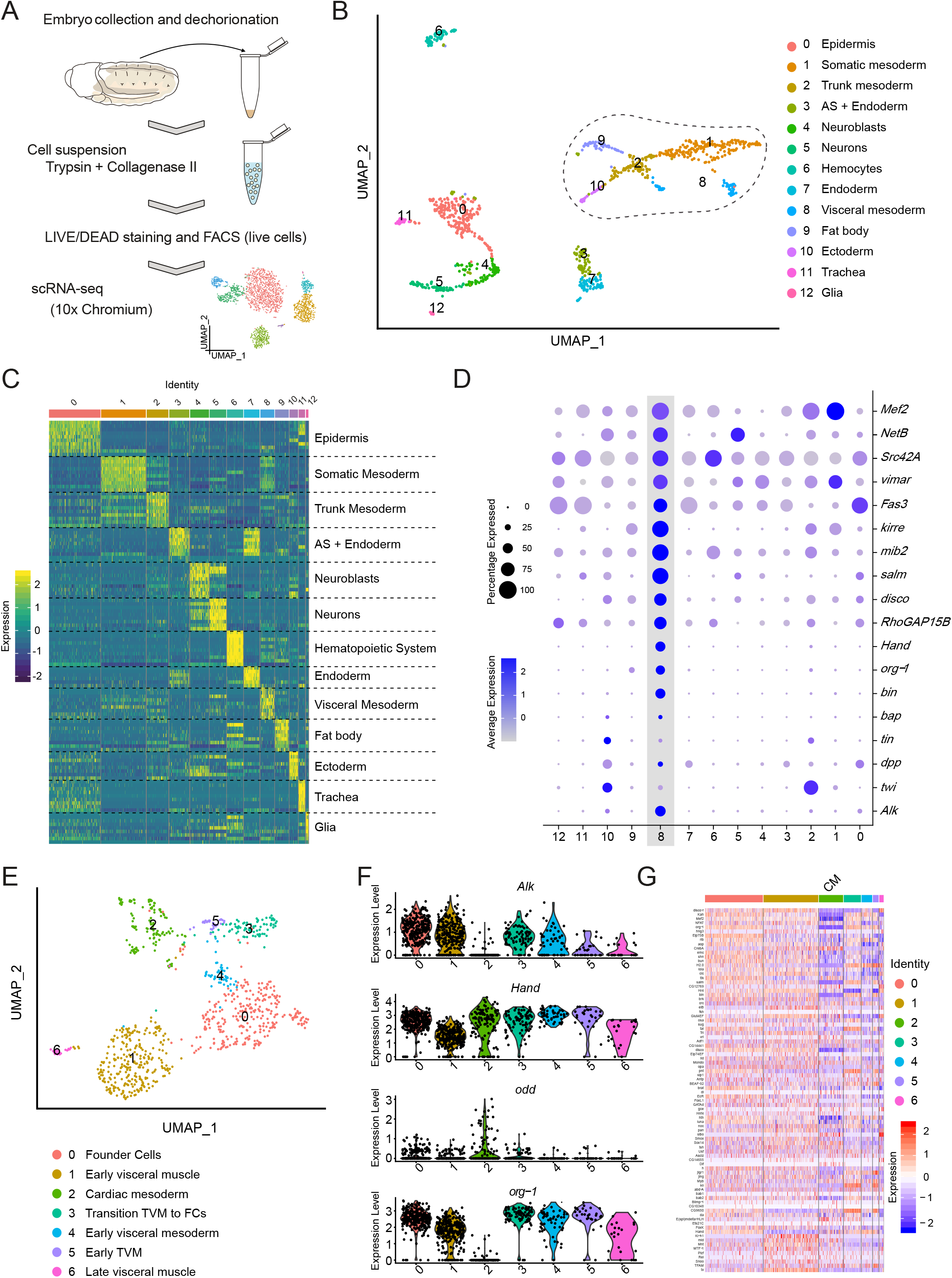
TaDa-identified Alk targets are enriched in the visceral mesoderm. **A.** Schematic outline of experimental approach. **B.** UMAP plot of whole embryo scRNA-seq showing 13 identified cell clusters arranged in four main groups. A dashed line surrounds the group formed by cell populations of mesodermal origin (1, 2, 8, 9 and 10). **C.** Heat map indicating the predicted identity of clusters 0-12, and their likely origin based on their gene expression profiles. **D.** Dot plot highlighting the increased expression of known factors involved in VM development, such as *bin, bap, org-1, Hand* and *Fas3,* in cluster 8. **E.** UMAP plot of *HandC-GFP* positive, FACS sorted cells revealing 7 individual cell clusters. **F.** Violin plots indicating expression of *Alk*, *Hand*, *odd* (expressed in the cardiac mesoderm) and *org-1* in clusters 0-7. **G.** Heat map indicating the relative expression of TaDa-identified targets downstream of Alk in clusters 0-7, highlighting an inverse correlation with cluster 2 (cardiac mesoderm).

We next wanted to analyze expression levels of the TaDa-identified candidates within the VM cluster. However, this was not possible for our whole embryo dataset due to (i) the overall low number of VM cells, and (ii) the low proportion of VM FCs that precluded a rigorous interrogation of TaDa candidate expression in relation to Alk activity. To achieve this, GFP-expressing cells were enriched using cell sorting by flow cytometry from *HandC-GFP; twi2xPE-Gal4>UAS-jeb* embryos with an enlarged visceral FC population. In this experiment, after quality filtering, we identified 888 cells. The cells distributed into seven clusters based on gene expression profiles, six of which exhibited VM identity (Fig. 4E). The remaining cluster (cluster 2) represents the *HandC-GFP-positive* cells of the cardiac mesoderm (CM), as indicated by their combinatorial expression of CM specific genes (Fig. 4F-G). A heatmap analysis revealed an enrichment of TaDa-identified Alk-regulated genes in the VM that was most prominent in clusters 0 and 1 (identified as FCs and early VM, respectively) when compared to the CM cluster (cluster 2)(Fig. 4G). Taken together, our single cell analysis supports the reliability of TaDa in identifying novel targets downstream of Alk signalling in the VM.

### *Kahuli* transcription is regulated by Alk signalling in the developing VM

Our TaDa analysis identified 151 TFs that are potentially regulated by Alk signalling activity (Fig. 2C-D) (Table S1). We chose to further investigate one of these: the Snail family TF Kahuli (Kah; Fig. 3B). The Snail TF family in *Drosophila* comprises Snail (Sna), together with Worniu (Wor) and Escargot (Esg) while the Scratch A and B families comprise Scratch (Scrt), CG12605 and Kah, that share a common domain structure with a Zinc finger C2H2-type DNA binding domain (Fig. 5A-A’). Kah is the only Scratch A family member and lacks the Scratch domain found in Scratch B family members (Kerner et al., 2009). *Kah* mRNA is expressed in a dynamic pattern, initially detected in early embryogenesis (Fig. S3A). Expression is seen at stage 9 in the developing VM, and later is enriched in FCs. We also found *Kah* expression in the somatic mesoderm (SM), where it is observed from stage 10 until late stage 16 (Fig. 5B, D). At the end of embryogenesis, *Kah* is expressed in the brain and ventral nerve cord (VNC) (Fig. S3A). Our *in situ* data identifying *Kah* expression in both the VM and SM was confirmed in our single cell RNA-seq datasets (Fig. 5B-C).

**Figure 5.**
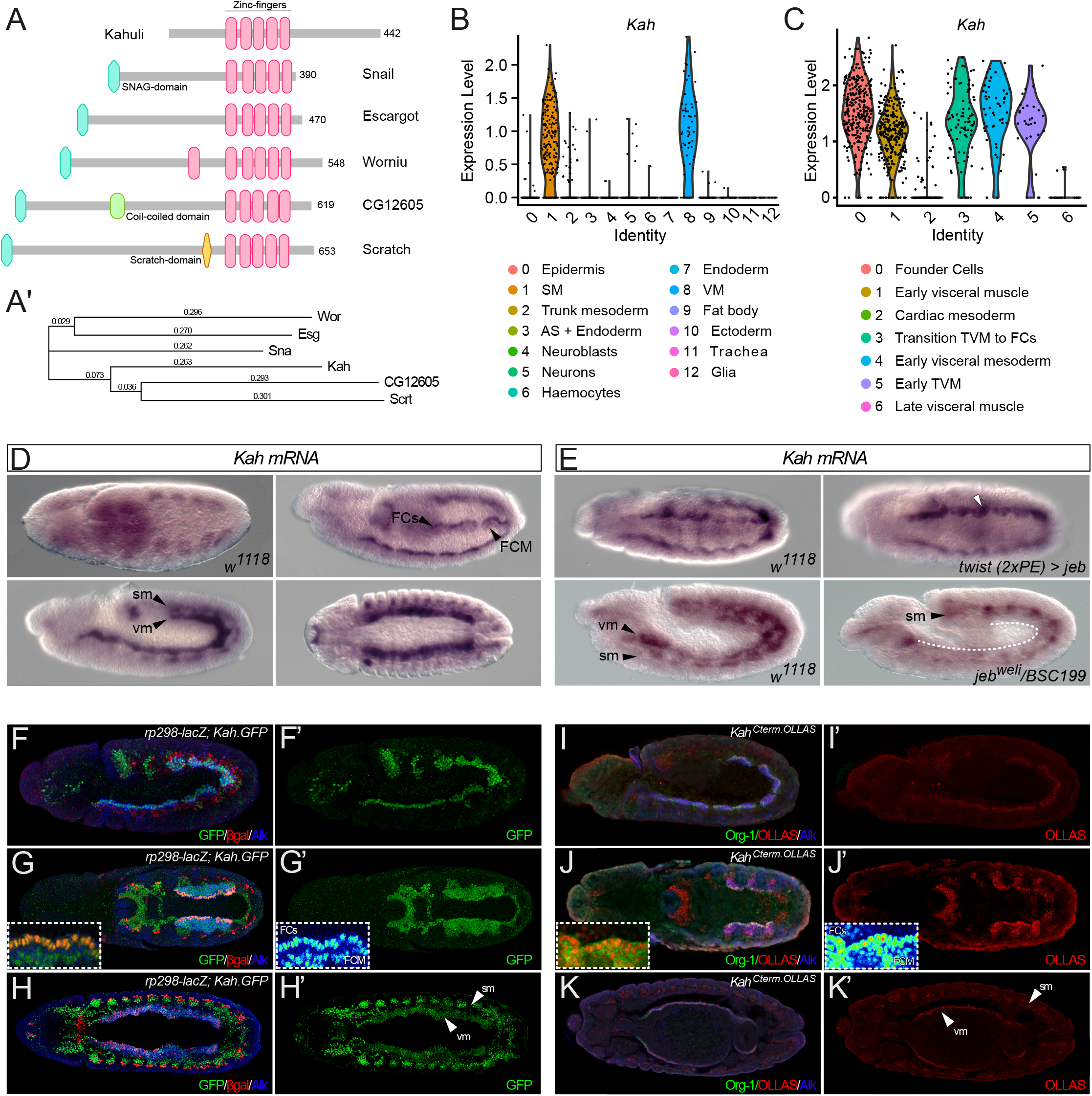
*Kahuli* is expressed in the developing visceral and somatic mesoderm. **A.** Kah belongs to the Snail/Scratch family of transcription factors sharing 5 zinc-finger domains. Schematic indicates the overall domain structure of the *Drosophila* family members, Kahuli, Snail, Escargot, Wornoi, CG12605 and Scratch. **A’.** Phylogenetic tree indicating the relationship of between Kah and the members of the Snail/Scratch family. **B.** Violin plots from scRNA-seq of wild-type embryos indicates *Kah* transcript is expressed in the embryonic VM and SM. **C.** Violin plots from scRNA-seq analysis of FACS sorted *Hand-GFP* expressing cells reveals expression of *Kah* mRNA in visceral, but not cardiac, mesoderm. **D.** *Kah* transcripts are abundant in SM and VM during embryogenesis, with increased expression levels in the visceral founder cell (FC) row. FCM, fusion competent myoblasts; sm, somatic musculature; vm, visceral musculature. **E.** Ectopic expression of *jeb* results in an increase of *Kah* expression in visceral FCMs. Conversely, animals devoid of Jeb/Alk signalling (*jeb^weli^* mutants) lack the strong FC-specific *Kah* expression in the VM while SM expression remains unaltered. **F-G’.** A Kah.GFP gene duplication construct can be detected from stage 10 embryos in the VM, with no clear distinction between FCs (marked by *rp298-lacZ,* red, inset depicts a close-up in LUT colors) and FCMs. Lateral view (F, F’), dorsal view (G, G’). **H-H’.** After myoblast fusion (stage 13), Kah.GFP is still maintained in visceral (vm) and somatic musculature (sm). Dorsal view. **I-J’.** Expression of endogenously-tagged Kah^Cterm.OLLAS^ is similar to Kah.GFP, but appears to be enriched in visceral FCs (marked by Org-1, green, inset depicts a close-up in LUT colors). Lateral view (I, I’), dorsal view (J, J’). **K-K’.** Stage 13 embryos show Kah^Cterm.OLLAS^ both in the visceral and somatic muscles (vm and sm, respectively). Dorsal view.

The robust *Kah* mRNA signal observed in visceral FCs was consistent with our hypothesis that *Kah* is a novel target of Alk activity. To test this further, we assessed *Kah* expression in the VM upon ectopic activation of Alk (on *jeb* overexpression) as well as in *jeb* mutants in which Alk is not activated. As predicted, ectopic Alk activation led to strong, uniform *Kah* expression throughout the entire VM, while loss of Alk signalling in *jeb* mutant embryos reduced *Kah* expression (Fig. 5E, Fig. S3B). As expected, *Kah* expression was still observed in the embryonic SM, as this tissue was unaffected by loss of Alk signalling (Fig. 5E, Fig. S3C). Thus, in agreement with the TaDa experimental approach, our experimental validation confirms that *Kah* is an Alk target gene in the VM.

### Kahuli protein is detected in the embryonic VM

To visualize the distribution of Kah protein during embryo development, we employed the BAC clone CH322-97G04 strain from the modERN (model organism Encyclopedia of Regulatory Networks), which carries an extra copy of the *Kah* locus encoding a C-terminally GFP::FLAG-tagged variant of Kah, expressed under control of endogenous regulatory elements (Kudron et al., 2018). This strain was generated by targeted genomic integration of the *Kah.GFP* recombining BAC into an intronic region of the *Msp300* gene and does not compromise fly viability (Fig. S4A). Notably, Kah.GFP can be detected throughout the VM and SM, in agreement with the expression of *Kah* mRNA (Fig. 5F-H’). However, in contrast with our mRNA analyses, high levels of Kah-GFP protein were observed in both VM FCs and FCMs (Fig. 5G-G’). Given the large size of GFP and its tendency to form homodimers at high concentrations (Yang et al., 1996) we were concerned that may impact function and stability of the Kah fusion protein, prompting us to generate a *Kah* allele with a C-terminal 3xOLLAS tag using CRISPR/Cas9 induced HDR (Fig. 5I-K’, Fig. 6A, Fig. S5; referred as *Kah^Cterm.OLLAS^).* Viable *Kah^Cterm.OLLAS^* animals were obtained that displayed nuclear OLLAS-tag staining in the visceral and somatic muscle. *Kah^Cterm.OLLAS^* was enriched in nuclei of FCs in the VM, in keeping with our *Kah* mRNA observations (Fig. 5J-J’). Taken together, these findings confirm that Kah is strongly expressed in the Alk-positive, developing VM and suggest that both Kah mRNA and protein levels are regulated by Alk signalling.

**Figure 6.**
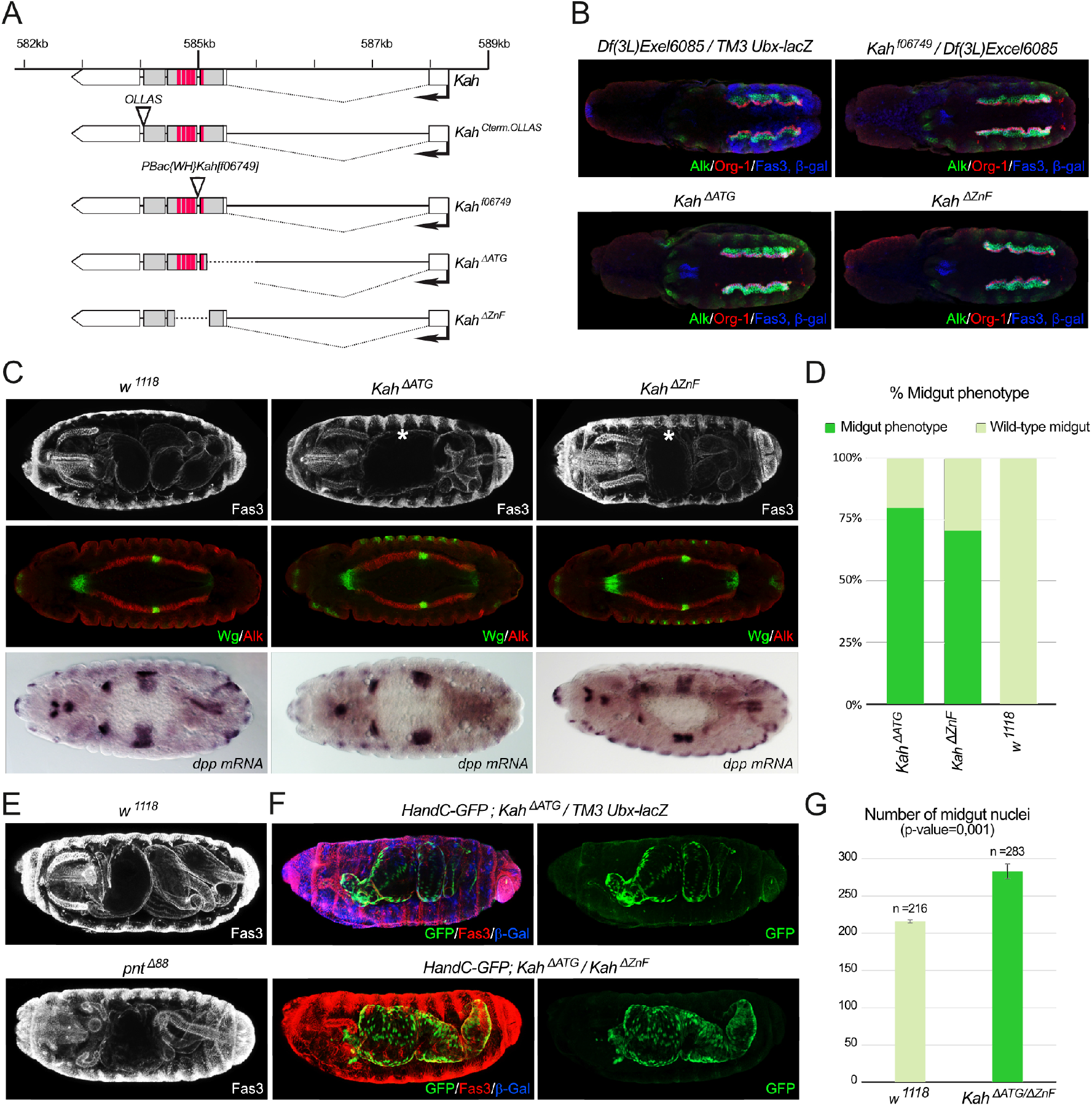
*Kah* mutants exhibit defects in midgut constriction. **A.** Schematic overview of the newly generated *Kah* alleles: *Kah^Cterm.OLLAS^, Kah^ΔATG^* and *Kah^ΔZnf^*, together with the *Kah^f06749^* PiggyBac insertion allele. **B.** Stage 10 embryos exhibit FC-specific expression of Org-1 (red) and *HandC-GFP* (green) FC markers. Dorsal views. **C.** Wild-type embryos at stage 16 are characterized by three midgut constrictions while *Kah* mutants fail to form the first midgut constriction. *Kah* mutants express Wg and *dpp* at levels comparable with control (*w^1118^*) embryos. Dorsal views. **D.** Quantification of the midgut constriction phenotype observed in C. **E.** *pnt^Δ88^* mutants display a midgut constriction phenotype similar to that observed in *Kah* mutants. Dorsal views. **F.** *Kah* mutants display abnormal organisation of midgut musculature, visualized with the nuclear *HandC-GFP* reporter FC markers (green). Lateral views. **G.** Quantification of the number of nuclei present in wild-type (*w^1118^*) and *Kah* mutants.

### Kahuli is required during embryonic midgut constriction

Given the expression of *Kah* in the VM, we next addressed a potential role of Kah in this tissue during embryonic development. We initially characterised the *Kah^f06749^* PiggyBac insertion and *Df(3L)Exel6085,* which deletes the entire *kah* locus (Fig. 6A, Fig. S4B’) in a lethality test. 38% of transheterozygous *Kah^f06749^/Df(3L)Exel6085* animals hatched, but 0% survived until L2 larvae (n=200). In our initial observations we did not identify any defects in FC specification, but instead noted abnormalities in the midgut of *Kah^f06749^/ Df(3L)Exel6085* and *Df(3L)Exel6085* embryos at later stages (Fig. 6B, Fig. S6). To further explore these findings, we generated two additional *Kah* alleles using CRISPR/Cas9; (i) *Kah^ΔATG^,* in which a deletion removes most of exon 2 including the predicted ATG start codon, and (ii) *Kah^ΔZnF^,* carrying an in-frame deletion that removes the region coding for the zinc-finger domains of Kah (Fig. 6A). Surprisingly, and in contrast to *Kah^f06749^, Kah^ΔATG^*and *Kah^ΔZnF^* were viable over *Df(3L)Exel6085* although we noticed a slightly increased lethality (75%, n=200 embryos) at the embryonic stage when compared to *w^1118^* controls. To investigate whether these *Kah* mutants exhibited defects in visceral cell fate specification or VM morphology, we visualized Alk, Fasciclin 3 (Fas3) and Org-1 at stage 11-12 in *Kah^f06749^/Df(3L)Exel6085, Kah^ΔATG^* and *Kah^ΔZnF^* embryos. For all *Kah* alleles we noted that Alk expression and localization was similar to controls (Fig. 6B). In addition, loss of Kah did not affect early VM cell identity, as indicated by FC-specific expression of Org-1 (Fig. 6B).

We next looked at later stages of gut development in *Kah^ΔATG^* and *Kah^AZnF^* mutants. After specification, FCs fuse with FCMs to form binucleate myotubes ultimately forming a web of interconnected muscles that surrounds the midgut endoderm (Georgias et al., 1997; Klapper et al., 2002; Kusch and Reuter, 1999; Martin et al., 2001; Rudolf et al., 2014). By stage 16, the midgut of wild-type embryos has acquired three constrictions that finally subdivide it into four chambers (Fig. 6C) (Campos-Ortega, 1997; Poulson, 1950; Reuter and Scott, 1990; Schroter et al., 2006). Using *HandC-GFP* and Fasciclin 3 (Fas3) to visualise the midgut, we found that the first midgut constriction in the newly generated *Kah^ΔATG^* and *Kah^ΔZnF^* mutants was frequently not formed or incomplete (*Kah^ΔATG^* 80%, n=89 embryos; *Kah^AZnF^* 70%, n=109 embryos) (Fig. 6C, asterisk; quantified in Fig. 6D). This highly penetrant phenotype was similar to that observed in *Kah^f06749^/Df(3L)Exel6085* as well as in *Df(3L)BSC362/Df(3L)Exel6085* embryos in which *Kah* is entirely deleted (Fig. S4B and S6A). Since Wg and Dpp signalling events are important for proper midgut constriction, we investigated their expression in *Kah* mutants. Both Wg protein and *dpp* mRNA levels were indistinguishable from controls, suggesting that the defective midgut constriction observed in the absence of *Kah* is not due to loss of Wg or Dpp (Fig. 6C).

We next turned towards transcription factors whose loss has been shown to affect VM constriction. Interestingly, the ETS TF Pointed (Pnt) exhibits defects in midgut constriction (Bilder et al., 1998), and a physical interaction with Kah has been reported by Y2H (Thurmond et al., 2019), http://flybi.hms.harvard.edu/results.php, prompting us to further investigate similarities between *pnt* and *Kah* mutants. Analysis of the amorphic *pnt^Δ88^* allele confirmed the previously described midgut constriction phenotype (Fig. 6E) (Bilder et al., 1998). Using the *HandC-GFP* reporter we revealed that *Kah* mutant embryos have an increased number of visceral nuclei in the midgut (Fig. 6F, G; n=283, p<0.001). These HandC-GFP nuclei were also highly disorganized when compared to controls where visceral muscle nuclei are aligned in four rows (Fig. 6F). We next performed epistasis experiments between our *Kah* loss of function alleles and *pnt^Δ88^.* When tested in a complementation analysis, we observed an increased lethality of transheterozygous *pnt^Δ88^/kah* loss of function animals. In addition, we found that approximately 23 % of late stage transheterozygous embryos exhibited defects in midgut constriction. Taken together, these findings suggest that Kah and Pnt could function together to effectively accomplish the first midgut constriction.

### RNA-seq identifies Kah target genes

Having identified a role for Kah in midgut constriction, we next performed RNA-seq on *Kah* mutants to identify putative targets of this previously uncharacterized TF. RNA-seq was performed on both *Kah^ΔATG^* and *Kah^ΔZnF^* mutants at embryonic stages 11-16. We noted 1664 and 2640 genes that were upregulated (log2FC ≥0.59 [FC≥1.5] and padj-value ≤0.05) while 1191 and 2827 genes were downregulated (log2FC ≤-0.59 [FC≤-1.5] and padj-value ≤0.05) in *Kah^ΔATG^* and *Kah^ΔZnF^* mutants, respectively (Fig. 7A-, Table S2). Further comparison identified 2524 overlapping genes differentially regulated (log2FC≥0.59 and ≤-0.59 [FC≥1.5 and ≤-1.5] at padj-value≤0.05) in both *Kah* mutants (1464 present at increased expression levels and 1034 at decreased expression levels), whose expression was correlated significantly (Fig. 7C-D). Many of the genes identified have yet to be investigated and represent interesting candidates for future functional characterization (Fig. 7E-F).

**Figure 7.**
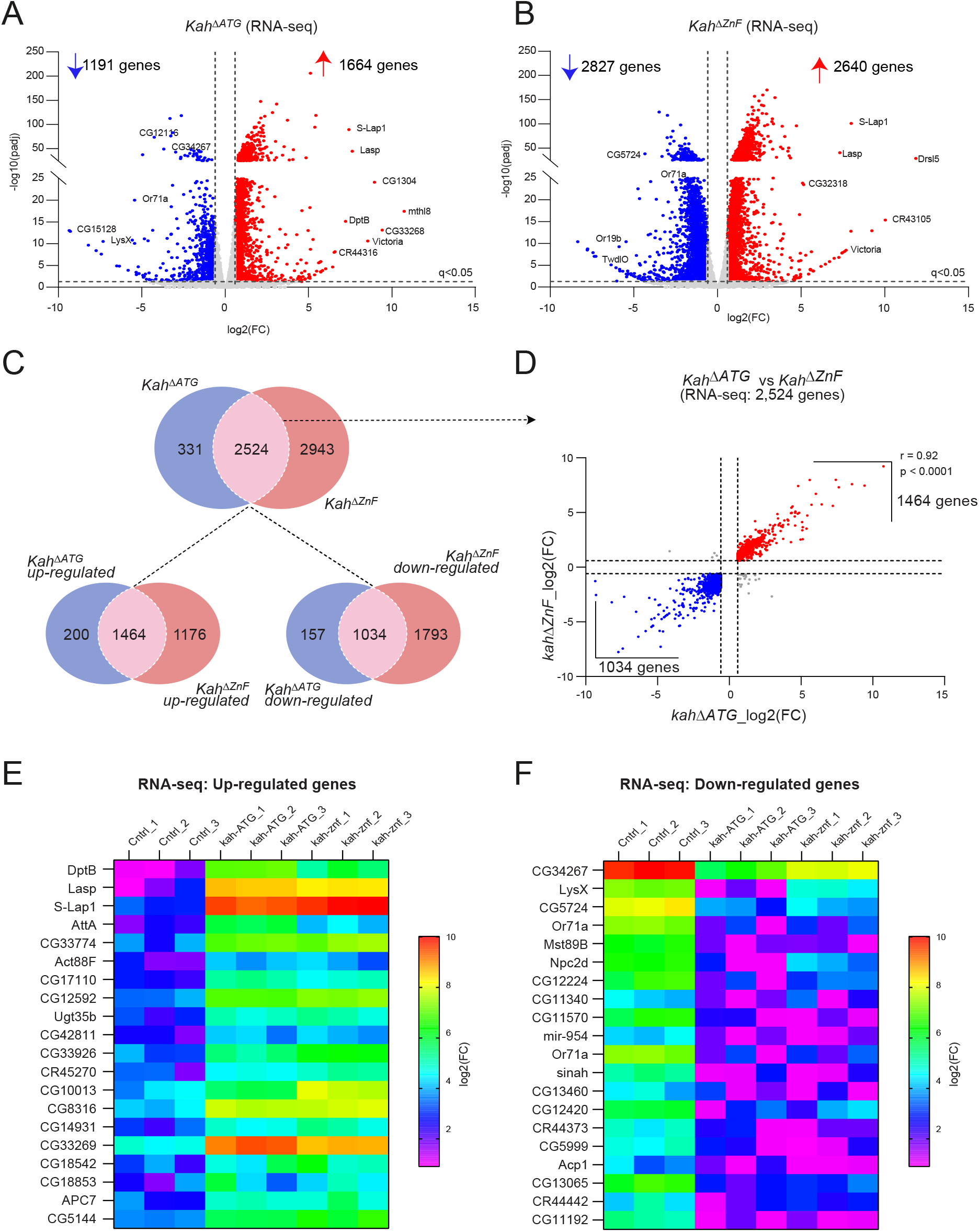
RNA-seq analysis identifies Kah target genes. **A-B.** Volcano plots of RNA-seq based differential gene expression measured in *Kah^ΔATG^* and *Kah^ΔZnF^* mutant embryos. See Table S2 for detailed results. Dashed lines show differential gene expression thresholds (Fold change ≥1.5 and ≤-1.5 (log2FC ≥0.59 and ≤-0.59) for up- and for down-regulated genes respectively (padj-value ≤0.05). Up-/down-regulated genes are indicated in red and blue respectively. A selection of genes that are differentially expressed are labeled. **C.** Venn diagrams indicating the number of differentially expressed genes observed in *Kah^ΔATG^* and *Kah^ΔZnF^* mutants. Top panel - all significantly differentially expressed genes; lower left panel - significantly differentially expressed upregulated genes, lower right panel - significantly differentially expressed downregulated genes. **D.** Correlation between the significantly differentially expressed genes (2,524) observed in *Kah^ΔATG^* and *Kah^ΔZnf^* mutants. Thresholds used to determine differential expression are indicated by dashed lines (Fold change ≥1.5 and ≤-1.5 (log2FC≥0.59 and ≤-0.59, and padj-value≤0.05). Pearson correlation coefficient is indicated at the top right of the plot. **E-F.** Differential gene expression heatmap of 20 highly differentially expressed genes (rows) in *Kah^ΔATG^* and *Kah^ΔZnf^* mutants compared with controls (Ctrl). Colour key is indicated adjacent: red - highest expression, blue - lowest expression.

### Analysis of Kah and Pnt ChIP-seq datasets identifies common targets and a Kah binding motif

ChIP-seq data performed in embryos with Kah-GFP has been deposited publicly by the modENCODE project (accession #ENCSR161YRO) (Roy et al., 2010). This Kah-GFP ChIP dataset contained a predominance of promoter regions with a peak in the vicinity of transcription start sites (TSS) (Fig. 8A, Fig. S7A). Basic Motif search in regions 50 bp and 200 bp region around the peaks led to the generation of a *de novo* motif for Kah that has highest scoring similarity with the related Snail TF (Fig. 8B). Among those genes containing a Kah motif in the vicinity of the TSS we noted a number expressed in the visceral mesoderm, such as *antennapedia (Antp), mind bomb 2 (mib2*) and *Netrin-b (NetB*) Table S3. To better evaluate Kah function at the level of gene expression, we compared the differentially expressed genes identified in *Kah* mutants by RNA-seq with the Kah ChIP-seq dataset. We found that 31% (339/1094) of genes identified in the Kah ChIP-seq dataset were differentially regulated in *Kah* mutant embryos (Fig. 8C, Table S3). We manually examined the publicly available *in situ* data (Thurmond et al., 2019), Berkeley *Drosophila* Genome Project (Hammonds et al., 2013; Tomancak et al., 2002; Tomancak et al., 2007), identifying 20% as having a curated expression in the embryonic midgut (Fig. 8D-E). Of note, many genes (42%) did not have annotated expression data in and could not be analysed.

**Figure 8.**
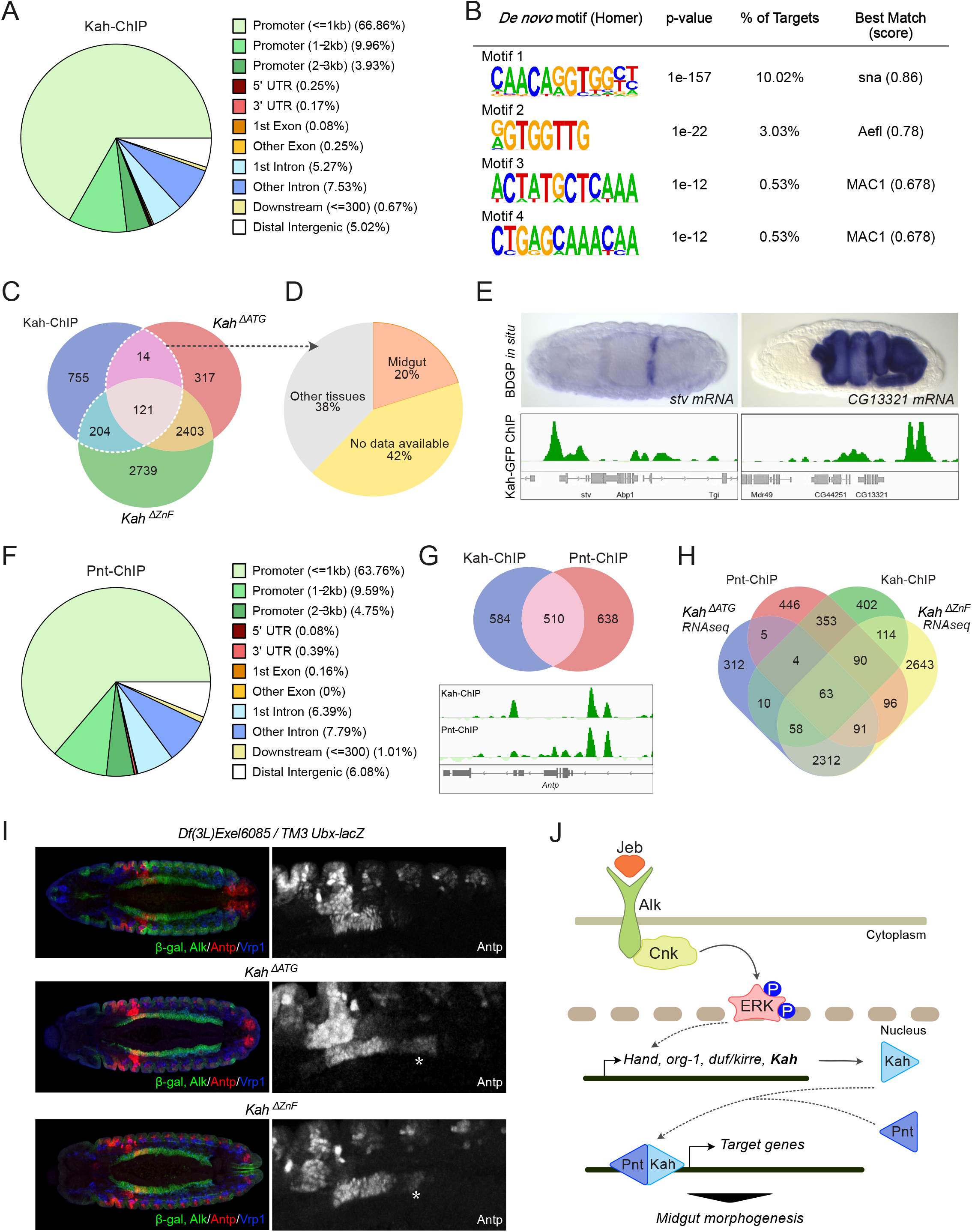
ChIP analysis identifies a *Kah* putative binding site and putative common targets of Kah and Pnt. **A.** Pie chart indicating Kah ChIP-seq peak locations in the genome, relative to promoter, UTR, intron/exon and other regions. **B.** Analysis of motif enrichments in regions of 50 bp around the peak center identifies a putative Kah binding motif highly related to the Sna binding motif. **C.** Venn diagram indicating the number of differentially expressed genes between *Kah^ΔATG^* and *Kah^ΔZnF^* mutants, compared with Kah-ChIP read location as indicated. **D.** Proportion of genes expressed in the midgut compared to other tissues or expression data not available. **E.** BDGP *in situ mRNA* expression pattern of *stv* and *CG13321* (differentially expressed in *Kah* mutants and Kah-ChIP targets) in the VM. Lower panel shows Kah ChIP peak profiles within the two candidate genes: *stv* and *CG13321.* **F.** Pie chart showing Pnt ChIP-seq peak locations in the genome, relative to promoter, UTR, intron/exon and other regions. **G.** Venn diagram showing the proportion of overlapping genes between Kah- and Pnt-ChIP datasets. Lower panel shows overlapping Pnt and Kah peaks within the *Antp* locus. **H.** Venn diagram showing the overlap between Kah-ChIP, Pnt-ChIP, and *Kah* mutant RNA-seq datasets. Details can be found in Table S2. **I.** *Kah^ΔATG^* and *Kah^ΔZnf^* display aberrant Antp protein expression in the visceral mesoderm. Embryos (stage 13) are stained for Antp (red), Alk (green), Vrp1 (blue) and β-gal (green) as indicated. **J.** Model for Alk regulation of Kah in the VM. Alk activation in the VM is driven by binding of its ligand Jeb, which induces the MAPK signalling cascade and activates the transcription of FC-specific genes including *Hand, org-1, duf/kirre* and the newly identified *Kah*. Kah may form a complex with other transcriptional regulators such as Pnt to target genes involved in the formation of the first midgut constriction.

ChIP-seq data from transgenic *Pnt-eGFP* embryos has also been deposited publicly by the modENCODE project (accession #ENCSR997UIM) (Roy et al., 2010), allowing us to compare Pnt and Kah binding locations throughout the genome, together with genes differentially expressed in *Kah* mutants. This analysis revealed that 46% (510/1094) of Kah-ChIP targets are potentially occupied by both Kah and Pnt, including *Antp* (Fig. 8G, Table S3). Further, 30% (157/510) of these common ChIP targets overlapped with genes differentially expressed in *Kah* mutants (Fig. 8H, Table S3). Since Antp is known to play an important role in setting up the first midgut constriction (Bilder et al., 1998; Roy et al., 1997), we examined Antp expression in *Kah* mutants, observing an expansion of Antp protein domain in the visceral mesoderm (Fig. 8I). Taken together, our analysis identifies a set of genes that are potentially regulated by Kah and Pnt downstream of Alk signalling during midgut constriction and worthy of further investigation in the future (Fig. 8J).

## Discussion

### Alk targets in the VM

Specification of muscle FCs in the visceral muscle primordia is dependent on Alk signalling in response to Jeb secretion from the somatic mesoderm (Englund et al., 2003; Lee et al., 2003; Stute et al., 2004). It is also known that signalling via Alk activates the Ras/MAPK pathway translocating the FCM fate-promoting transcription factor Lameduck (Lmd) from the nucleus to the cytoplasm (Popichenko et al., 2013). A similar mechanism has been suggested for a still unknown FC-fate repressor triggering the FC-specific transcriptional program in the VM (Popichenko et al., 2013; Zhou et al., 2019). This transcriptional program remains relatively unexplored with only a few identified targets reported, such as *Hand, org-1, kirre, dpp* and *Alk* itself (Englund et al., 2003; Lee et al., 2003; Mendoza-Garcia et al., 2017; Shirinian et al., 2007; Varshney and Palmer, 2006). Chromatin immunoprecipitation (ChIP) has been the predominant approach for mapping protein-chromatin interactions. However, ChIP assays require a great amount of starting material and a specific antibody, which is the main limitation of this technique (Wu et al., 2016). Furthermore, when investigating transcriptional regulatory proteins downstream of RTKs such as Alk, a broad range of TFs are likely to be involved precluding individual ChIP analyses. A second approach, RNA sequencing (RNA-seq), has also been intensely employed for transcriptomic analyses. While straight forward for cell culture studies, dissection or isolation of the tissue of interest, in this case the VM, would be required for its use in identifying Alk transcriptional targets in *Drosophila.* Therefore, in our efforts to identify novel transcriptional targets of Alk activity in the VM we employed a third option, the TaDa approach which allows genome-wide RNA Pol II occupancy to be investigated in the specific tissue of choice (Southall et al., 2013).

### TaDa reproduces endogenous RNA Pol II binding

Our experimental design was based on two conditions manipulating Alk signalling, one resulting in activation, and the other in inhibition, of Alk signalling throughout the VM, followed by TaDa analysis. Comparison of our TaDa dataset with previously published RNA-seq data (NCBI BioProject, accession number PRJEB11879) from cells isolated from the mesoderm suggest our data recapitulated endogenous binding of RNA Pol II. Our dataset also agreed with current understanding of Alk signalling and induction of cell fate specification in the trunk VM by activation of a FC-specific transcriptional program (Lee et al., 2003; Loren et al., 2003; Stute et al., 2004), including observed differential expression of previously identified Alk transcriptional targets such as *Hand, org-1, kirre* and *dpp.* Taken together, a combination of different analyses supported our approach as replicating transcriptional events in the VM and led us to validate of TaDa-identified genes in the VM as targets of Alk-driven signalling events.

### TaDa identified VM-specific genes

A number of differentially expressed genes were validated by *in situ* hybridization during embryogenesis. For all selected genes, mRNA was indeed visualized in the visceral musculature. Expression of the membrane protein Fax was observed in the VM and the CNS, as previously reported (Hill et al., 1995). Interestingly, Fax has been identified in a screen for diet-regulated proteins in the *Drosophila* ovary (Hsu and Drummond-Barbosa, 2017). Insulin signalling in response to diet promotes activation of the Ribosomal protein S6 Kinase (S6K), that drives *fax* expression, resulting in the extension of ovarian niche escort cell membranes the via cytoskeleton remodeling (Hsu and Drummond-Barbosa, 2017; Su et al., 2018). Notably, Alk has been reported to modulate insulin signalling in the brain during nutrient restriction, via activation of the common downstream target PI3-kinase (Cheng et al., 2011; Okamoto and Nishimura, 2015), making Fax an interesting candidate for further study.

Another interesting candidate is Pdp1, which has been reported to have differentially expressed mRNA isoforms through the use of multiple enhancers and promoters (Reddy et al., 2000). Our *in situ* probe was designed to detect all six isoforms; therefore, we are unable to identify which Pdp1 isoform is expressed in the VM from our present analysis. Moreover, Pdp1 has been reported to function as either a transcriptional activator or repressor, depending on the isoform. Interestingly, Pdp1 has been reported to play an important role during muscle formation by promoting expression of *Tropomyosin I* (*TmI*) (Reddy et al., 2000). Identification of the specific isoform expressed in the VM will be important to further address its function downstream of Alk in the VM (Lin et al., 1997). Our TaDa analysis identified numerous genes that have not been characterized to date, and further investigation will be crucial to decipher their role *in vivo* in particular during the formation of the visceral circular muscles.

### *Kahuli* plays a role in later visceral musculature development

One of the interesting uncharacterized targets was *Kah* (*Kahuli*), which encodes a Snail family transcription factor. Overexpression of Kah in the thorax has been reported to block development of thoracic bristles, revealing a potential to drive changes in cell identity (Singari et al., 2014). Alk signalling in the VM imposes a change in the transcriptional program, leading to FC specification (Englund et al., 2003; Lee et al., 2003; Stute et al., 2004). We were able to validate *Kah* as an Alk target locus, with clear differences in *Kah* expression when Alk signalling was either blocked or activated. However, we also noted Alk-independent *Kah* transcription in the early VM, similar to that already described for *org-1* (Schaub and Frasch, 2013), in addition to the Alk modulated transcription. Currently, the transcription factors downstream of Alk that regulate *Kah* transcription are unknown, although this will be interesting to study in the future.

The role of Alk signalling in the VM is the differentiation of the ventral-most row of cells from the ingressed trunk VM into FCs (Englund et al., 2003; Lee et al., 2003; Stute et al., 2004). Alk-mediated FC specification occurs via activation of the Ras/MAPK pathway, leading to the transcription of FC-specific genes such as *Hand, org-1, kirre, dpp* and also *Kah* (Englund et al., 2003; Lee et al., 2003; Shirinian et al., 2007; Stute et al., 2004; Varshney and Palmer, 2006). Neither the loss of *Hand* nor *org-1* alters FC specification in the VM, likely reflecting a highly regulated process with redundant roles that assure FC and fusion of visceral muscles (Schaub and Frasch, 2013; Varshney and Palmer, 2006). Our characterization of *Kah* mutant alleles allow us to conclude that, similarly to *Hand* and *org-1,* Kah activity in the VM is dispensable for FC specification, although formally Kah could be responsible for FC specific transcriptional changes of yet unidentified targets.

*Kah* mutants are viable but exhibit defects in the process of midgut constriction formation. Work from a number of groups has implicated a group of players in this event, such as Wg, Dpp and Ubx as well as Pnt, Emc and Org-1 ((Bilder et al., 1998; Ellis et al., 1990; Muller et al., 1989; Panganiban et al., 1990; Reuter et al., 1990; Schaub and Frasch, 2013). Interestingly, Alk signalling activity is important for the expression of Dpp in the VM and the maintenance of *org-1* expression in FCs (Popichenko et al., 2013; Shirinian et al., 2007). In this study, we were particularly interested in Pnt, as this ETS domain TF has been reported by the FlyBi project (http://flybi.hms.harvard.edu/) to bind to Kah in high-throughput Y2H (Thurmond et al., 2019) and also to exhibit a midgut constriction phenotype (Bilder et al., 1998). Further, like Kah, Pnt is not required for FC specification in the VM (Zhou et al., 2019). We confirmed the midgut constriction phenotype in *pnt* mutants, which was similar to those observed in our *Kah* mutants. One obvious difference is in that the *pnt* mutant phenotype is completely penetrant, in comparison to that observed in *Kah* mutants. These data, together with the observation of a similar phenotype in transheterozygous *pnt^Δ88^/kah* embryos, suggest that Kah and Pnt may function together in transcriptional regulation of midgut constriction. Employing publicly available Kah-ChIP datasets (Roy et al., 2010) we were able to define a Kah binding motif, which was most similar to that described of the Snail TF. Further analysis of publicly available Pnt-ChIP datasets (Roy et al., 2010) highlighted *Antp* as a target of both Kah and Pnt binding during embryogenesis. *Antp* has a known role in gene expression in the VM and *Antp* mutants exhibit defects in midgut constriction (Bilder et al., 1998; Roy et al., 1997). Interestingly, while Antp appears to be misregulated in *Kah* mutants, Wg and *dpp* expression appears normal. Indeed, earlier work has reported that Wg and *dpp* expression are also normal in the VM of *pnt* mutant embryos (Bilder et al., 1998). Likely, as yet unidentified players function downstream of Kah in this process. Our bioinformatics analysis of *Kah* mutant RNA-seq datasets together with the Kah-ChIP and Pnt-ChIP datasets has identified a group of genes as candidates to be focused on in future studies, hopefully allowing us to better understand this process. It is also important to note that while in this work we have focused on a role in the VM, Kah is also expressed in the embryonic SM and CNS. As noted above, the differentially regulated genes identified in our *Kah* mutant RNA-seq analysis could be regulated by Kah in any of these tissues. Clearly, further investigation is needed to characterize the role of the Kah TF in the SM and CNS.

### Conclusions

Taken together, the use of the TaDa approach successfully allowed us to identify transcriptional targets of Alk signalling in the developing mesoderm, including the here novel target Kahuli described here. Many of these targets are currently uncharacterized and future studies should allow their function(s) in the VM to be elucidated. Our more in-depth study of Kah highlights a role for the TF in later visceral musculature development, where it appears to works in concert with other factors, including Pnt, to drive midgut constriction. Combined ChIP and RNA-seq analyses highlights a group of interesting, and largely uncharacterized genes, which should shed light on the midgut constriction process further.

## Materials and methods

### *Drosophila* stocks and genetics

LacZ or GFP balancer chromosomes were used to distinguish progeny of crosses. The following stock lines were obtained from the Bloomington *Drosophila* Stock Center (BDSC): *TM3 Sb Ubx-lacZ* (#9120), *Df(3L)BSC362 (Kah* deficiency, #24386) *Df(3L)Exel8065 (Kah* deficiency, #7564), *Kah^f06749^* (PiggyBac insertion mapped to the second intron of *Kah,* #19006), *Kah-GFP.FPTB* (#64829), *twi.2xPE-Gal4* (#2517), *Df(2R)BSC199 (jeb* deficiency, #9626). Additional stocks used in this study: *UAS-LT3-NDam-Pol II* (Southall et al., 2013), *rP298-lacZ* (Nose et al., 1998) (an enhancer trap in the *kirre* locus (Ruiz-Gomez et al., 2000)), *HandC-GFP* (Sellin et al., 2006), *bap3-Gal4* (Zaffran et al., 2001), *UAS-jeb.V* (Varshney and Palmer, 2006), *UAS-Alk.EC.MYC* (Bazigou et al., 2007) (Alk extracellular domain that functions as dominant negative, here referred to as *UAS-Alk.DN*), *jeb^weli^* (Stute et al., 2004), *Alk^10^* (Loren et al., 2003).

### TaDa sample preparation

*Twi.2xPE-Gal4* and *bap3-Gal4* lines were used to test Dam-Pol II toxicity and further RNA-Pol II profiling. Embryos were collected over a 4 h period and aged at 25°C to stage 10-13, followed by dechorionation in 2% hydroxychloride solution for 2 min and subsequent washing steps in PBS. A total of 50 μl embryos per sample was used as starting material. Genomic DNA was extracted (QIAGEN Blood and Tissue DNA extraction kit) and methylated DNA processed and amplified as previously described (Choksi et al., 2006; Sun et al., 2003), with the following modifications. Upon verification of non-sheared gDNA, the DpnI digestion was set up in 50 μl. After overnight DpnI digestion, the DNA was purified (QIAGEN MinElute PCR purification kit) into 30 μl of MQ water, from where 15 μl were used for the adaptors-ligation step. Amplified DNA from experimental and Dam-only embryos was again purified (QIAGEN MinElute PCR purification kit) into 20 μl of MQ water and 200 ng aliquots were run in a 1% agarose gel to verify amplification of different fragments (visualized as a smear from 500 bp to 2-3 kb). Purified PCR products were used for PCR-free library preparation, followed by pair-end sequencing on Illumina HiSeq X Ten platform (BGI Tech Solutions, Hong Kong). Three biological replicates were performed for transcriptional profiling of the visceral mesoderm on each of the experimental genetic backgrounds.

### DamID-seq bioinformatics data analysis

The *Drosophila* genome (FASTA) and genes (GTF) version r6.21 were downloaded from Flybase (Gramates et al., 2017; Thurmond et al., 2019) and all GATC regions extracted in BED format using fuzznuc (Rice et al., 2000). The paired FASTQ files from 18 samples (background Dam, Jeb, and DN at three replicates each for *twi.2xPE-Gal4* and *bap3-Gal4*) were aligned to the *Drosophila* genome using *bowtie2 (--very-sensitive-local*) (Langmead and Salzberg, 2012). *Sambamba (merge*) (Tarasov et al., 2015) was used to combine replicates and the log-fold changes between DN/Jeb and Dam, obtained using *bamCompare (--centerReads --normalizeTo1x 142573017 --smoothLength 5 -bs 1*) from deepTools (Ramirez et al., 2014). Counts of reads mapped on edge to GATC fragments were generated using a script (*GATC_mapper.pl*) from DamID-Seq pipeline (Maksimov et al., 2016). RNAseq data from a public dataset (PRJEB11879) was used to quantify the expression of genes (only 6-8h mesoderm samples used). The GATC level counts were converted to gene level counts using intersectBed from Bedtools (Quinlan, 2014) and compared against the gene expression (only background Dam samples) at TPM level. GATC sites were merged into peaks following the methods prescribed in a previous study (Tosti et al., 2018). In brief logFC for individual GATCs were generated using *Limma* (Jeb vs Dam and DN vs Dam) (P < 1e-5) and the GATC sites were merged into peaks based on median GATC fragment distance in the *Drosophila* genome using *mergeWindows* and *combineTests function* from the *csaw* package (Lun and Smyth, 2016). The peaks were assigned to overlapping genes and filtered for FDR at 0.01. The final results were taken only for the DN vs Dam comparison. The Jeb samples had low rate of alignment, hence Jeb vs Dam is only used as a visual confirmation of the DN vs Dam peaks at specific locations. Enrichment of transcription factors in the peaks generated was performed by using Fisher test against list of published *Drosophila* transcription factors (Kudron et al., 2018). Enrichment for GO and KEGG terms for the genes assigned to significant peaks was performed using *WebGestalt* (Liao et al., 2019). All statistical analysis was performed in the R programming environment.

### Embryo dissociation into single cells and cell sorting

Embryos were collected on apple juice agar plates and aged to stage 10-13 (as confirmed by microscopic visualization of a small fraction). Embryos were dechorionated 2% hydorxychloride solution for 2 min and washed in cold PBS. Subsequently, embryos were incubated in dissociating solution (1 mg/ml trypsin, 0.5 collagenase I, 2 % BSA) for 1 h and vortexed every 10 minutes, after which the reaction was stopped by addition of 10 volumes of ice cold PBS. Dissociated cell solution was sieved through 70 μm and 40 μm cell strainers to remove cell clumps. Dead cells or debris from the dissociated samples were removed using the EasySep Dead Cell Removal (Anexin V) Kit (STEMCELL, ref. 17899) according to the manufacturer’s guidelines. The remaining cells were respectively labelled with aqua-fluorescent reactive dye (dying cells) and calcein violet AM (living cells) using the LIVE/DEAD Violet Viability/Vitality Kit (Molecular Probes, ref. L34958) under manufacturer’s guidelines. Finally, each sample was washed twice in PBS, 2% fetal bovine serum and resuspended in 500 μl PBS, 2% fetal bovine serum. Living cells were enriched using a FACSAria III cell sorter (BD biosciences) based on the LIVE/DEAD staining and, when appropriate, GFP expression driven by the Hand-GFP construct. The cells were sorted using an 85 μm nozzle into Eppendorf tubes that had been pre-coated with PBS/2% BSA.

### Generation of single cell libraries, sequencing and bioinformatic analysis

Approximately 2500 sorted cells were loaded onto one lane of a Chromium 10X chip (10X Genomics) and libraries prepared using the normal workflow for Single Cell 3’ v3 libraries (10X Genomics). Libraries were sequenced on the NextSeq 500 platform (Illumina), and the raw format base call files (BCLs) sequences were demuliplexed using cellranger mkfastq version 3.1. After read QC, mapping was performed with the *Drosophila Melanogaster* genome using STAR aligner. For analysis, unique molecular identified (UMI) count matrix were imported into the Seurat R toolkit version 3.1. For quality filtering, cells with less than 1000 genes and more than 5000 expressed genes were excluded. Also, cells which are expressed more than 25% mitochondrial genes removed. Subsequent count normalization, scaling, feature selection, clustering (PCA) and dimensionality reduction (UMAP-Uniform Manifold Approximation and Projection) were performed according to the standard workflow (Stuart et al., 2019).

### CRISPR/Cas9 mediated generation of mutant and tagged *Kah* alleles

Deletion (*Kah^ΔATG^* and *Kah^ΔZnF^*) and endogenous tagged (*Kah^Cterm.OLLAS^) Kah* alleles were generated using the CRISPR/Cas9 system. CRISPR target sites were identified and evaluated using flyCRISPR Optimal Target Finder tool (Gratz et al., 2015). Single guide RNA (sgRNA) target sequences (sequences available in Table S4) were cloned into pU6-BbsI-chiRNA vector (Addgene, Cat. No. 45946) and injected into *vasa-Cas9* (BDSC, #51323) embryos (BestGene Inc.). For *Kah^Cterm.OLLAS^* a donor construct was added to the injection mix. The injected animals were crossed to third chromosome balancer flies (BDSC, #9120) and their progeny were PCR-screened for positives deletion/insertion events. Positive candidates were confirmed further by Sanger sequencing (Eurofins Genomics).

Endogenously tagged *Kah^Cterm.OLLAS^* was generated using CRISPR/Cas9 induced homology directed repair (HDR) at the *Kah* c-terminal. Three CRISPR sgRNA sequences (Sequences available in Table S4) were used. One sgRNA was designed to target upstream to the *Kah* stop codon and the other two to target directly the *Kah* stop codon. In addition, a DNA donor cassette was synthesized (Integrated DNA Technologies, Inc.), prior to Gibson assembly cloning into the pBluescript II KS[-] HDR plasmid. This donor cassette codes for the remaining part of the *Kah* c-terminal followed by six glycines (linker), three copies of the OLLAS-tag and a TAG stop codon; flanked by two homology arms (495bp upstream and 500bp downstream of the respective Cas9 cutting sites, with codon optimized target sequences).

### ChIP bioinformatics based determination of Kah and Pnt binding motifs

Publicly available *dm6* aligned Kah ChIP-seq data input libraries (modEncodeID: ENCSR161YRO & ENCSR664RUV) and the raw format FASTQ sequences of Pnt Chip-sequences were retrieved from (modEncodeID: ENCSR997UIM & ENCSR249WKC). For Pnt-Chip seq data, the base quality of each sequenced read was assessed using the FASTQC program. The reads were aligned to the *Drosophila melanogaster* (BDGP6) reference genome using Bowtie2. Due to the ambiguity of reads that align to multiple locations across the genome, only reads that uniquely mapped were considered for subsequent analysis. Post alignment processes were performed with samtools and BEDtools, and Homer suite v4.1 program “findpeaks” (Heinz et al., 2010) with default transcription factor finding parameters (-style factor) used for peak calling. Resulting peaks from each replicate were annotated using ChIPseeker v1.2 (Yu et al., 2015) and merged to be used as input for genome-wide motif enrichment scanning using the “findMotifsGenome.pl” script from the Homer suite. Regions of 50 bp and 200 bp around the peak center were analyzed for motif enrichment.

### ChIP-seq visualization

BigWig files were used to search for DNA occupancy (Peak profile) for respective genes on Integrative Genomics Viewer (version 2.8.12).

### RNA-sequencing and analysis

Embryos were collected at 25°C (5-16 h after egg laying) and dechorionated in 2% hydroxychloride solution for 4 min. After subsequent washing with embryo wash (0.8% NaCl and 0.05% Triton X-100) and H20, embryos were stored at −80°C. One embryo collection for 3 biological replicates per genotype was obtained, RNA-extraction was carried out according to the manufacturer’s protocol (Promega ReliaPrepTM RNA Tissue Miniprep System, REF-Z6111). Total RNA was measured using NanoDrop OneC (Thermo Scientific) for its concentration and RNA-integrity was checked on the gel electrophoresis. 9-15 μg of total RNA/biological replicate was shipped to Novogene Co. Ltd (UK) for sequencing. Prior to making library, samples were reassessed for quality with the Agilent 2100 Bioanalyzer system. Sequencing was performed on an Illumina platform and paired-end reads were produced. Over 40 million reads/genotype were generated and mapped to the genome at a rate of over 96%. Drosophila melanogaster (ensamble bdgp6_gca_000001215_4 genome assembly) was used. HISAT2 algorithm for alignment and DESeq2 R package (Anders and Huber, 2010) for differential gene expression was used. Subsequent analyses was performed with Microsoft Excel 2016 and Graphpad Prism 9. Fold change ≥1.5 and ≤-1.5 (log2FC≥0.59 and ≤-0.59) for up- and down regulated genes respectively, and padj-value≤0.05 was used for statistical significance.

### Immunohistochemistry

Embryos were fixated and stained as described (Müller, 2008). Primary antibodies used were: guinea pig anti-Alk (1:1000 (Loren et al., 2003)), rabbit anti-Alk (1:750 (Loren et al., 2003)), chicken anti-β-galactosidase (1:200; Abcam ab9361), mouse anti-Fasciclin3 (1:50; DSHB 7G10), mouse anti-Antp (1:50; DSHB 4C3) rabbit anti-GFP (1:500; Abcam ab290), chicken anti-GFP (1:300; Abcam ab13970), mouse anti-Wg (1:50, DSHB 4D4), rat anti-OLLAS (1:200, pre-absorbed on *w^1118^* embryos; Abnova), rabbit anti-Org-1 (1:1000, (Mendoza-Garcia et al., 2017)), sheep anti-digoxygenin-AP fab fragment 1:4000 (Roche). Alexa Fluor®-conjugated secondary antibodies were from Jackson Immuno Research. Embryos were dehydrated in an ascending ethanol series before clearing and mounting in methylsalicylate. Images were acquired with a Zeiss LSM800 confocal microscope or Axiocam 503 camera, processed and analyzed employing Zeiss ZEN2 (Blue Edition) imaging software. Nuclei quantification (Fig. 6G) was performed with ImageJ software (Schneider et al., 2012). Raw images were converted into binary format and nuclei were quantified with 3D nuclei counter package (Bolte and Cordelieres, 2006).

### *In situ* hybridization

For in situ hybridization, fragments of the respective CDS were PCR amplified from genomic DNA, cloned into the dual promoter pCRII-TA vector (ThermoFisher, #K207040) and used as template to generate DIG-labeled in situ probes with SP6/T7 polymerases (Roche, # 10999644001). Whole-mount *in situ* hybridization was done according to Lécuyer et al. (Lecuyer et al., 2008), with modifications adapted from (Pfeifer et al., 2012). Samples were mounted using *in situ* mounting media (Electron Microscopy Sciences). Images were acquired with a Zeiss Axio Imager.Z2 microscope, processed and analyzed employing Zeiss ZEN2 (Blue Edition) imaging software.

## Acknowledgments

We thank A. Brand, M. Takeichi, M. Frasch, A. Paululat, A. Nose, M. Ruiz-Gomez and J. Bateman for sharing fly stocks and reagents, and B. Paul and C. Schaub for valuable help and suggestions. We acknowledge Bloomington *Drosophila* Stock Center (NIH P40OD018537) for fly stocks used in this study. The Fasciclin3, Wg and Antp antibodies developed by C. Goodman, S. M. Cohen and D. Brower, respectively, were obtained from the Developmental Studies Hybridoma Bank, created by the NICHD of the NIH and maintained at The University of Iowa, Department of Biology, Iowa City, IA 52242. This work has been supported by grants from the Swedish Cancer Society (RHP CAN18/0729; EL CAN15/541; JLA CAN17/342), the Children’s Cancer Foundation (RHP 2019-0078), the Swedish Research Council (RHP 2019-03914; EL 14-3596; JLA 2016-03306; MB 24400813), the Swedish Foundation for Strategic Research (RB13-0204), the Göran Gustafsson Foundation (RHP2016) and the Knut and Alice Wallenberg Foundation KAW 2015.0144).

## Author contributions

S.B., V.A., H.L., J.L., P.M.G. and E.L. carried out the bioinformatics analysis. P.M.G., S.K.S., B.A., L.M., G.W. and M.B. performed the wet lab experiments. R.H.P. supervised the project. R.H.P. and P.M.G. wrote the first manuscript draft that was further developed with all authors.

## References

Anders, S. and Huber, W. (2010). Differential expression analysis for sequence count data. Genome Biol 11, R106.

Aughey, G. N. and Southall, T. D. (2016). Dam it’s good! DamID profiling of protein-DNA interactions. Wiley Interdiscip Rev Dev Biol 5, 25–37.

Bazigou, E., Apitz, H., Johansson, J., Loren, C. E., Hirst, E. M., Chen, P. L., Palmer, R. H. and Salecker, I. (2007). Anterograde Jelly belly and Alk receptor tyrosine kinase signaling mediates retinal axon targeting in Drosophila. Cell 128, 961–975.

Bilder, D., Graba, Y. and Scott, M. P. (1998). Wnt and TGFbeta signals subdivide the AbdA Hox domain during Drosophila mesoderm patterning. Development 125, 1781–1790.

Bolte, S. and Cordelieres, F. P. (2006). A guided tour into subcellular colocalization analysis in light microscopy. J Microsc 224, 213–232.

Brand, A. H. and Perrimon, N. (1993). Targeted gene expression as a means of altering cell fates and generating dominant phenotypes. Development 118, 401–415.

Campos-Ortega, J. A. a. H., V (1997). The Embryonic Development of Drosophila melanogaster. Berlin, Germany: Springer.

Cheng, L. Y., Bailey, A. P., Leevers, S. J., Ragan, T. J., Driscoll, P. C. and Gould, A. P. (2011). Anaplastic lymphoma kinase spares organ growth during nutrient restriction in Drosophila. Cell 146, 435–447.

Choksi, S. P., Southall, T. D., Bossing, T., Edoff, K., de Wit, E., Fischer, B. E., van Steensel, B., Micklem, G. and Brand, A. H. (2006). Prospero acts as a binary switch between self-renewal and differentiation in Drosophila neural stem cells. Dev Cell 11, 775–789.

Ellis, H. M., Spann, D. R. and Posakony, J. W. (1990). extramacrochaetae, a negative regulator of sensory organ development in Drosophila, defines a new class of helix-loop-helix proteins. Cell 61, 27–38.

Englund, C., Loren, C. E., Grabbe, C., Varshney, G. K., Deleuil, F., Hallberg, B. and Palmer, R. H. (2003). Jeb signals through the Alk receptor tyrosine kinase to drive visceral muscle fusion. Nature 425, 512–516.

Frasch, M. (1995). Induction of visceral and cardiac mesoderm by ectodermal Dpp in the early Drosophila embryo. Nature 374, 464–467.

Georgias, C., Wasser, M. and Hinz, U. (1997). A basic-helix-loop-helix protein expressed in precursors of Drosophila longitudinal visceral muscles. Mech Dev 69, 115–124.

Gramates, L. S., Marygold, S. J., Santos, G. D., Urbano, J. M., Antonazzo, G., Matthews, B. B., Rey, A. J., Tabone, C. J., Crosby, M. A., Emmert, D. B., et al. (2017). FlyBase at 25: looking to the future. Nucleic Acids Res 45, D663–D671.

Gratz, S. J., Rubinstein, C. D., Harrison, M. M., Wildonger, J. and O’Connor-Giles, K. M. (2015). CRISPR-Cas9 Genome Editing in Drosophila. Curr Protoc Mol Biol 111, 31 32 31–31 32 20.

Hammonds, A. S., Bristow, C. A., Fisher, W. W., Weiszmann, R., Wu, S., Hartenstein, V., Kellis, M., Yu, B., Frise, E. and Celniker, S. E. (2013). Spatial expression of transcription factors in Drosophila embryonic organ development. Genome Biol 14, R140.

Heinz, S., Benner, C., Spann, N., Bertolino, E., Lin, Y. C., Laslo, P., Cheng, J. X., Murre, C., Singh, H. and Glass, C. K. (2010). Simple combinations of lineage-determining transcription factors prime cis-regulatory elements required for macrophage and B cell identities. Mol Cell 38, 576–589.

Hill, K. K., Bedian, V., Juang, J. L. and Hoffmann, F. M. (1995). Genetic interactions between the Drosophila Abelson (Abl) tyrosine kinase and failed axon connections (fax), a novel protein in axon bundles. Genetics 141, 595–606.

Hsu, H. J. and Drummond-Barbosa, D. (2017). A visual screen for diet-regulated proteins in the Drosophila ovary using GFP protein trap lines. Gene Expr Patterns 23-24, 13–21.

Kerner, P., Hung, J., Behague, J., Le Gouar, M., Balavoine, G. and Vervoort, M. (2009). Insights into the evolution of the snail superfamily from metazoan wide molecular phylogenies and expression data in annelids. BMC Evol Biol 9, 94.

Klapper, R., Stute, C., Schomaker, O., Strasser, T., Janning, W., Renkawitz-Pohl, R. and Holz, A. (2002). The formation of syncytia within the visceral musculature of the Drosophila midgut is dependent on duf, sns and mbc. Mech Dev 110, 85–96.

Kudron, M. M., Victorsen, A., Gevirtzman, L., Hillier, L. W., Fisher, W. W., Vafeados, D., Kirkey, M., Hammonds, A. S., Gersch, J., Ammouri, H., et al. (2018). The ModERN Resource: Genome-Wide Binding Profiles for Hundreds of Drosophila and Caenorhabditis elegans Transcription Factors. Genetics 208, 937–949.

Kusch, T. and Reuter, R. (1999). Functions for Drosophila brachyenteron and forkhead in mesoderm specification and cell signalling. Development 126, 3991–4003.

Langmead, B. and Salzberg, S. L. (2012). Fast gapped-read alignment with Bowtie 2. Nat Methods 9, 357–359.

Lecuyer, E., Parthasarathy, N. and Krause, H. M. (2008). Fluorescent in situ hybridization protocols in Drosophila embryos and tissues. Methods in molecular biology (Clifton, N.J 420, 289–302.

Lee, H.-H., Zaffran, S. and Frasch, M. (2006). Development of the Larval Visceral Musculature. In Muscle Development in Drosophila, pp. 62–78. New York, NY: Springer New York.

Lee, H. H., Norris, A., Weiss, J. B. and Frasch, M. (2003). Jelly belly protein activates the receptor tyrosine kinase Alk to specify visceral muscle pioneers. Nature 425, 507–512.

Liao, Y., Wang, J., Jaehnig, E. J., Shi, Z. and Zhang, B. (2019). WebGestalt 2019: gene set analysis toolkit with revamped UIs and APIs. Nucleic Acids Res 47, W199–W205.

Lin, S. C., Lin, M. H., Horvath, P., Reddy, K. L. and Storti, R. V. (1997). PDP1, a novel Drosophila PAR domain bZIP transcription factor expressed in developing mesoderm, endoderm and ectoderm, is a transcriptional regulator of somatic muscle genes. Development 124, 4685–4696.

Loren, C. E., Englund, C., Grabbe, C., Hallberg, B., Hunter, T. and Palmer, R. H. (2003). A crucial role for the Anaplastic lymphoma kinase receptor tyrosine kinase in gut development in Drosophila melanogaster. EMBO Rep 4, 781–786.

Loren, C. E., Scully, A., Grabbe, C., Edeen, P. T., Thomas, J., McKeown, M., Hunter, T. and Palmer, R. H. (2001). Identification and characterization of DAlk: a novel Drosophila melanogaster RTK which drives ERK activation in vivo. Genes Cells 6, 531–544.

Lun, A. T. and Smyth, G. K. (2016). csaw: a Bioconductor package for differential binding analysis of ChIP-seq data using sliding windows. Nucleic Acids Res 44, e45.

Maksimov, D. A., Laktionov, P. P. and Belyakin, S. N. (2016). Data analysis algorithm for DamID-seq profiling of chromatin proteins in Drosophila melanogaster. Chromosome Res 24, 481–494.

Martin, B. S., Ruiz-Gomez, M., Landgraf, M. and Bate, M. (2001). A distinct set of founders and fusion-competent myoblasts make visceral muscles in the Drosophila embryo. Development 128, 3331–3338.

Mendoza-Garcia, P., Hugosson, F., Fallah, M., Higgins, M. L., Iwasaki, Y., Pfeifer, K., Wolfstetter, G., Varshney, G., Popichenko, D., Gergen, J. P., et al. (2017). The Zic family homologue Odd-paired regulates Alk expression in Drosophila. PLoS Genet 13, e1006617.

Muller, J., Thuringer, F., Biggin, M., Zust, B. and Bienz, M. (1989). Coordinate action of a proximal homeoprotein binding site and a distal sequence confers the Ultrabithorax expression pattern in the visceral mesoderm. EMBO J 8, 4143–4151.

Müller, H. A. J. (2008). Immunolabelling of embryos. Methods in molecular biology (Clifton, N.J.) 420, 207–218.

Nose, A., Isshiki, T. and Takeichi, M. (1998). Regional specification of muscle progenitors in Drosophila: the role of the msh homeobox gene. Development 125, 215–223.

Okamoto, N. and Nishimura, T. (2015). Signaling from Glia and Cholinergic Neurons Controls Nutrient-Dependent Production of an Insulin-like Peptide for Drosophila Body Growth. Dev Cell 35, 295–310.

Panganiban, G. E., Reuter, R., Scott, M. P. and Hoffmann, F. M. (1990). A Drosophila growth factor homolog, decapentaplegic, regulates homeotic gene expression within and across germ layers during midgut morphogenesis. Development 110, 1041–1050.

Pfeifer, K., Dorresteijn, A. W. and Frobius, A. C. (2012). Activation of Hox genes during caudal regeneration of the polychaete annelid Platynereis dumerilii. Dev Genes Evol 222, 165–179.

Popichenko, D., Hugosson, F., Sjogren, C., Dogru, M., Yamazaki, Y., Wolfstetter, G., Schonherr, C., Fallah, M., Hallberg, B., Nguyen, H., et al. (2013). Jeb/Alk signalling regulates the Lame duck GLI family transcription factor in the Drosophila visceral mesoderm. Development 140, 3156–3166.

Poulson, D. F. (1950). Histogenesis, organogenesis and differentiation in the embryo of Drosophila melanogaster Meigen. Biology of Drosophila., 168–274.

Quinlan, A. R. (2014). BEDTools: The Swiss-Army Tool for Genome Feature Analysis. Curr Protoc Bioinformatics 47, 11 12 11–34.

Ramirez, F., Dundar, F., Diehl, S., Gruning, B. A. and Manke, T. (2014). deepTools: a flexible platform for exploring deep-sequencing data. Nucleic Acids Res 42, W187–191.

Reddy, K. L., Wohlwill, A., Dzitoeva, S., Lin, M. H., Holbrook, S. and Storti, R. V. (2000). The Drosophila PAR domain protein 1 (Pdp1) gene encodes multiple differentially expressed mRNAs and proteins through the use of multiple enhancers and promoters. Dev Biol 224, 401–414.

Reuter, R., Panganiban, G. E., Hoffmann, F. M. and Scott, M. P. (1990). Homeotic genes regulate the spatial expression of putative growth factors in the visceral mesoderm of Drosophila embryos. Development 110, 1031–1040.

Reuter, R. and Scott, M. P. (1990). Expression and function of the homoeotic genes Antennapedia and Sex combs reduced in the embryonic midgut of Drosophila. Development 109, 289–303.

Rice, P., Longden, I. and Bleasby, A. (2000). EMBOSS: the European Molecular Biology Open Software Suite. Trends Genet 16, 276–277.

Roy, S., Ernst, J., Kharchenko, P. V., Kheradpour, P., Negre, N., Eaton, M. L., Landolin, J. M., Bristow, C. A., Ma, L., Lin, M. F., et al. (2010). Identification of functional elements and regulatory circuits by Drosophila modENCODE. Science 330, 1787–1797.

Roy, S., Shashidhara, L. S. and VijayRaghavan, K. (1997). Muscles in the Drosophila second thoracic segment are patterned independently of autonomous homeotic gene function. Curr Biol 7, 222–227.

Rudolf, A., Buttgereit, D., Jacobs, M., Wolfstetter, G., Kesper, D., Putz, M., Berger, S., Renkawitz-Pohl, R., Holz, A. and Onel, S. F. (2014). Distinct genetic programs guide Drosophila circular and longitudinal visceral myoblast fusion. BMC Cell Biol 15, 27.

Ruiz-Gomez, M., Coutts, N., Price, A., Taylor, M. V. and Bate, M. (2000). Drosophila dumbfounded: a myoblast attractant essential for fusion. Cell 102, 189–198.

Satija, R., Farrell, J. A., Gennert, D., Schier, A. F. and Regev, A. (2015). Spatial reconstruction of single-cell gene expression data. Nat Biotechnol 33, 495–502.

Schaub, C. and Frasch, M. (2013). Org-1 is required for the diversification of circular visceral muscle founder cells and normal midgut morphogenesis. Dev Biol 376, 245–259.

Schneider, C. A., Rasband, W. S. and Eliceiri, K. W. (2012). NIH Image to ImageJ: 25 years of image analysis. Nat Methods 9, 671–675.

Schroter, R. H., Buttgereit, D., Beck, L., Holz, A. and Renkawitz-Pohl, R. (2006). Blown fuse regulates stretching and outgrowth but not myoblast fusion of the circular visceral muscles in Drosophila. Differentiation 74, 608–621.

Sellin, J., Albrecht, S., Kolsch, V. and Paululat, A. (2006). Dynamics of heart differentiation, visualized utilizing heart enhancer elements of the Drosophila melanogaster bHLH transcription factor Hand. Gene Expr Patterns 6, 360–375.

Shirinian, M., Varshney, G., Loren, C. E., Grabbe, C. and Palmer, R. H. (2007). Drosophila Anaplastic Lymphoma Kinase regulates Dpp signalling in the developing embryonic gut. Differentiation 75, 418–426.

Singari, S., Javeed, N., Tardi, N. J., Marada, S., Carlson, J. C., Kirk, S., Thorn, J. M. and Edwards, K. A. (2014). Inducible protein traps with dominant phenotypes for functional analysis of the Drosophila genome. Genetics 196, 91–105.

Southall, T. D., Gold, K. S., Egger, B., Davidson, C. M., Caygill, E. E., Marshall, O. J. and Brand, A. H. (2013). Cell-type-specific profiling of gene expression and chromatin binding without cell isolation: assaying RNA Pol II occupancy in neural stem cells. Dev Cell 26, 101–112.

Stuart, T., Butler, A., Hoffman, P., Hafemeister, C., Papalexi, E., Mauck, W. M., 3rd, Hao, Y., Stoeckius, M., Smibert, P. and Satija, R. (2019). Comprehensive Integration of Single-Cell Data. Cell 177, 1888–1902 e1821.

Stute, C., Schimmelpfeng, K., Renkawitz-Pohl, R., Palmer, R. H. and Holz, A. (2004). Myoblast determination in the somatic and visceral mesoderm depends on Notch signalling as well as on milliways(mili(Alk)) as receptor for Jeb signalling. Development 131, 743–754.

Su, Y. H., Rastegri, E., Kao, S. H., Lai, C. M., Lin, K. Y., Liao, H. Y., Wang, M. H. and Hsu, H. J. (2018). Diet regulates membrane extension and survival of niche escort cells for germline homeostasis via insulin signaling. Development 145.

Sun, L. V., Chen, L., Greil, F., Negre, N., Li, T. R., Cavalli, G., Zhao, H., Van Steensel, B. and White, K. P. (2003). Protein-DNA interaction mapping using genomic tiling path microarrays in Drosophila. Proc Natl Acad Sci U S A 100, 9428–9433.

Tarasov, A., Vilella, A. J., Cuppen, E., Nijman, I. J. and Prins, P. (2015). Sambamba: fast processing of NGS alignment formats. Bioinformatics (Oxford, England) 31, 2032–2034.

Thurmond, J., Goodman, J. L., Strelets, V. B., Attrill, H., Gramates, L. S., Marygold, S. J., Matthews, B. B., Millburn, G., Antonazzo, G., Trovisco, V., et al. (2019). FlyBase 2.0: the next generation. Nucleic Acids Res 47, D759–D765.

Tomancak, P., Beaton, A., Weiszmann, R., Kwan, E., Shu, S., Lewis, S. E., Richards, S., Ashburner, M., Hartenstein, V., Celniker, S. E., et al. (2002). Systematic determination of patterns of gene expression during Drosophila embryogenesis. Genome Biol 3, RESEARCH0088.

Tomancak, P., Berman, B. P., Beaton, A., Weiszmann, R., Kwan, E., Hartenstein, V., Celniker, S. E. and Rubin, G. M. (2007). Global analysis of patterns of gene expression during Drosophila embryogenesis. Genome Biol 8, R145.

Tosti, L., Ashmore, J., Tan, B. S. N., Carbone, B., Mistri, T. K., Wilson, V., Tomlinson, S. R. and Kaji, K. (2018). Mapping transcription factor occupancy using minimal numbers of cells in vitro and in vivo. Genome Res 28, 592–605.

Varshney, G. K. and Palmer, R. H. (2006). The bHLH transcription factor Hand is regulated by Alk in the Drosophila embryonic gut. Biochem Biophys Res Commun 351, 839–846.

Wolfstetter, G., Pfeifer, K., van Dijk, J. R., Hugosson, F., Lu, X. and Palmer, R. H. (2017). The scaffolding protein Cnk binds to the receptor tyrosine kinase Alk to promote visceral founder cell specification in Drosophila. Science signaling 10.

Wu, F., Olson, B. G. and Yao, J. (2016). DamID-seq: Genome-wide Mapping of Protein-DNA Interactions by High Throughput Sequencing of Adenine-methylated DNA Fragments. J Vis Exp, e53620.

Yang, F., Moss, L. G. and Phillips, G. N., Jr. (1996). The molecular structure of green fluorescent protein. Nat Biotechnol 14, 1246–1251.

Yu, G., Wang, L. G. and He, Q. Y. (2015). ChIPseeker: an R/Bioconductor package for ChIP peak annotation, comparison and visualization. Bioinformatics (Oxford, England) 31, 2382–2383.

Zaffran, S., Kuchler, A., Lee, H. H. and Frasch, M. (2001). biniou (FoxF), a central component in a regulatory network controlling visceral mesoderm development and midgut morphogenesis in Drosophila. Genes Dev 15, 2900–2915.

Zhou, Y., Popadowski, S. E., Deutschman, E. and Halfon, M. S. (2019). Distinct roles and requirements for Ras pathway signaling in visceral versus somatic muscle founder specification. Development 146.

